# The eukaryotic replisome intrinsically generates asymmetric daughter chromatin fibers

**DOI:** 10.1101/2025.09.18.677126

**Authors:** Bruna V Eckhardt, Hannah J Richter, Inge Rondeel, Keerthi Renduchintala, Francesca Mattiroli, Vijay Ramani

## Abstract

DNA replication is molecularly asymmetric, due to distinct mechanisms for lagging and leading strand DNA synthesis. Whether chromatin assembly on newly replicated strands is also asymmetric remains unknown, as visualizing this short-lived state in cells is impossible. To circumvent this limitation, we combine *in vitro* reconstitution of the *Saccharomyces cerevisiae* DNA and chromatin replication machineries with replication-aware single-molecule chromatin footprinting, to study how chromatin is re-assembled on replicated DNA. Leveraging the non-destructive, single-molecule, and strand-specific nature of our data, we discover an intrinsic asymmetry in nucleosome positioning patterns and organization between lagging- and leading-strand chromatin created by the yeast replisome. This asymmetry is only partially restored upon addition of chromatin assembly factors involved in *de novo* histone deposition and the ATP-dependent chromatin remodeler Isw1a, implying that other regulatory factors must resolve this asymmetry in cells. In sum, our data reveal the complexity of chromatin re-establishment following DNA replication, and suggest an asymmetric chromatin assembly intermediate on each daughter chromatid. These pathways have implications for essential chromatin-templated processes such as DNA repair, transcription, and gene silencing at replication forks.

## INTRODUCTION

During DNA replication, chromatin structure is accurately inherited to regulate cell fate after cell division (*1–4*). The complex responsible for DNA replication (*i*.*e*. the replisome) semi-conservatively synthesizes nascent lagging- and leading-strand DNA in an asymmetric fashion (*5*). A suite of histone chaperones – both distinct from and intrinsic to the replisome – then assemble chromatin onto nascent DNA, via a combination of parental histone recycling and *de novo* histone deposition (*6–25*). Importantly, this nascent chromatin state is critical to epigenetic memory, is short-lived, and has generally been thought to be assembled symmetrically (*1, 4*). This is because nucleosome modifications (*26*) and nucleosome unwrapping (*27*) appear to be equivalently apportioned according to most available measurements. This model, however, has not been rigorously tested, owing to i.) the inability to reliably isolate nascent chromatin from cells during this short-lived period; and, ii.) the inherent limitations of bulk averaging of chromatin structure from templates digested with destructive enzymes such as micrococcal nuclease (MNase) or the hyperactive transposase Tn5.

A solution to the first limitation can be found in the complete *in vitro* reconstitution of the *S. cerevisiae* replisome (*28–30*), which has revolutionized our understanding of the mechanisms of replication of free DNA, thanks to the ability to precisely control the process and characterize short-lived intermediates. Bottom-up reconstitutions have also illustrated that replication of chromatin templates can be recapitulated *in vitro* (*31, 32*) paving the way for new studies of nascent chromatin assembly. We recently presented a solution to the second limitation: a long-read sequencing technology that allows the identification of both DNA replication status and chromatin organization at single molecule level (*27, 33, 34*). However, due to the complexity of DNA and chromatin replication in cells, we did not achieve strand-specific resolution in this context.

Uniting these two approaches, we present here a new method that combines comprehensive *in vitro* reconstitutions of chromatin replication with replication-aware single-molecule chromatin footprinting. Our approach reveals strand-specific single-molecule nascent chromatin organization for the first time. By perturbing parental histone recycling, *de novo* histone deposition, and ATP-dependent nucleosome remodeling, we uncover that leading- and lagging-strands are organized into distinct chromatin fiber structures by replication forks, uncovering a structural ‘asymmetry’ between strands. This asymmetry is only partially resolved by addition of the *de novo* histone deposition pathway and remodeling factors, indicating that nascent chromatin assembles via an unanticipated strand-asymmetric intermediate state. These findings have crucial implications for understanding how daughter genomes and epigenomes are accurately inherited, and how replication-associated stress mechanisms may act.

## RESULTS

### Biochemical reconstitution of yeast chromatin replication and *de novo* chromatin assembly

We purified the components of the *S. cerevisiae* replisome to recapitulate all salient aspects of DNA replication (*i*.*e*. origin loading, licensing, firing, DNA replication elongation, lagging strand processing, and termination)(*28– 31*). This reconstitution included histone chaperones involved in parental histone recycling, as well as the *de novo* chromatin assembly pathway (32 preparations, 73 polypeptide chains; **Figure 1A**). As input, we used ∼10.5 kb chromatinized templates containing the origin of replication *ARS1*, prepared via Nap1/Isw1a-assisted assembly(*35*) in the presence of ORC to facilitate origin recognition within chromatin (**Figure 1B**)(*31*). Biochemical validation confirmed the functionality of our reconstitution, demonstrating both leading- and lagging-strand synthesis, their processing and termination in a DDK- and FACT-dependent manner (**Figure 1C**), and the formation of nucleosomes on these templates by MNase digestion (**Figure 1D**). Addition of the *de novo* chromatin assembly factors further enhances nucleosome formation in these reactions (**Figure 1E**), confirming its successful integration into this recombinant system. These results attest to our ability to reconstitute DNA replication, replisome-mediated parental histone recycling, and *de novo* nucleosome assembly during DNA synthesis.

**Figure 1:**
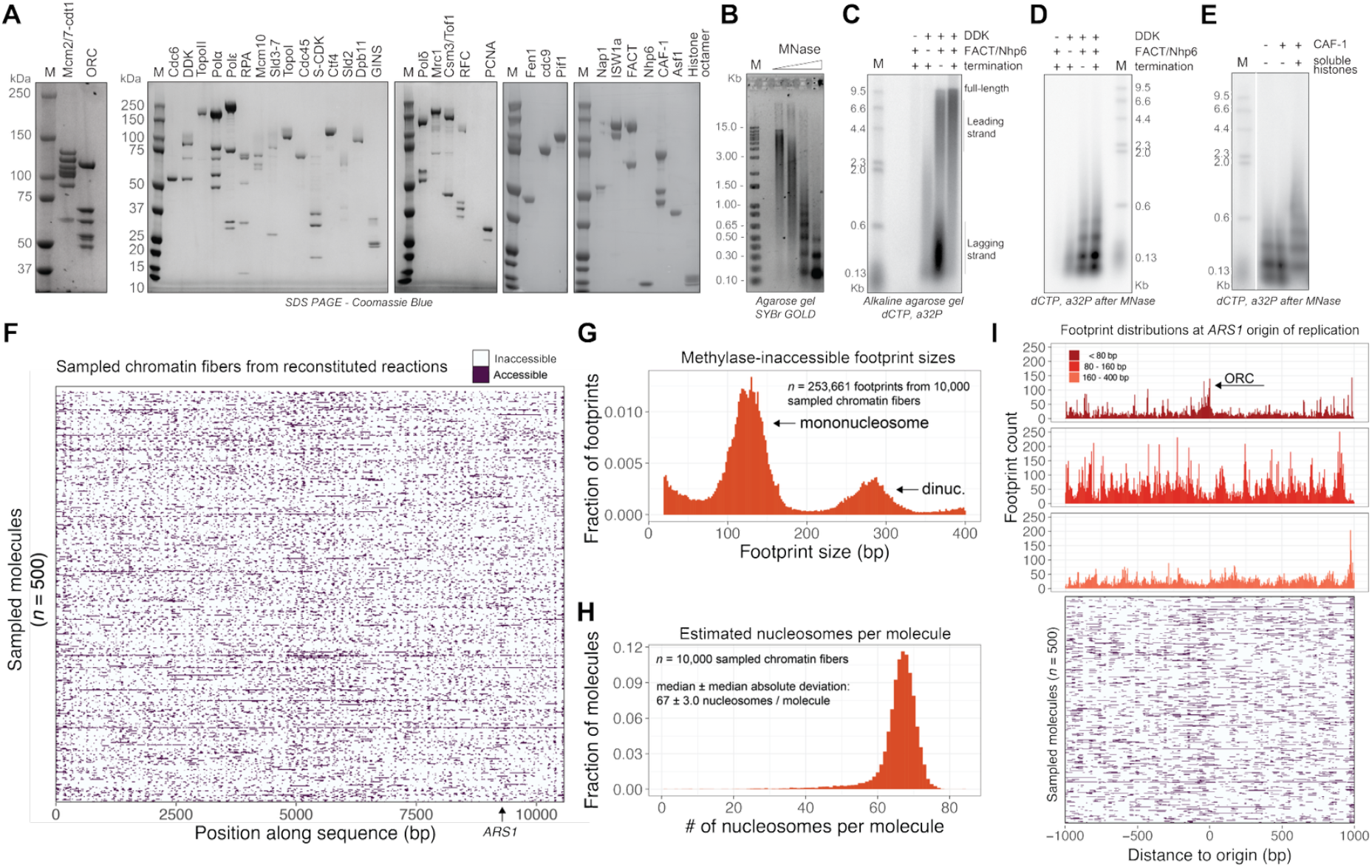
Single-molecule footprinting of reconstituted chromatin replication reactions. **(A)** SDS-PAGE of purified *S. cerevisiae* factors used to reconstitute chromatin replication. **(B)** SybrGOLD staining of a native agarose gel after Micrococcal Nuclease (MNase) digestion of input chromatin, demonstrating protected DNA fragments, indicative of chromatin assembly on a 10.5 kb template DNA containing yeast origin *ARS1*. **(C)** Autoradiography scan of a denaturing agarose gel of DNA replication products from chromatin replication reactions, confirming DDK and FACT dependency of our chromatin replication reactions. Fen1, Cdc9 and Pif1 (termination mix) are also active and process replicated DNA. **(D)** Autoradiography scan of a native agarose gel after MNase digestion of the samples shown in panel C, confirms chromatin assembly on replicated DNA (parental histone recycling). **(E)** Autoradiography scan of a native agarose gel after MNase digestion of reactions containing *de novo* chromatin assembly components. **(F)** Heatmap representation of 500 sampled, footprinted molecules, where each line represents a single molecule, purple represents predicted methyltransferase accessible stretches of DNA, and off-white represents methyltransferase-inaccessible stretches of DNA (as defined by a neural network – hidden Markov model). **(G)** Methyltransferase-inaccessible footprint size distributions for 10,000 sampled chromatin fibers, with expected mononucleosome and dinucleosome footprint sizes annotated. **(H)** Gaussian mixture model-based estimation of number of nucleosomes per footprinted molecule for 10,000 sampled molecules, with median and median absolute deviation annotated inset. **(I)** Size-stratified footprint midpoint position analyses focused on the *ARS1* origin of replication with short (< 80 bp; dark red), nucleosomal (80 – 160 bp; medium red), and oligonucleosomal (160 + bp; light red) footprint midpoint frequencies tabulated surrounding the *ARS1* center. Bottom: Heatmap visualization of 500 sampled single-molecules focusing on the same window.

### Single-molecule footprinting of reconstituted chromatin fiber replication reactions

To establish a readout to monitor chromatin organization in these reactions, we used SAMOSA to test <u>Ch</u>romatin <u>A</u>ccessibility on <u>A</u>ssembled <u>T</u>emplates (SAMOSA-ChAAT) (*33*), which uses single-molecule long-read chromatin footprinting by the m^6^dA methyltransferase EcoGII (*36*) to characterize biochemically reconstituted chromatin templates (schematic in **Supplementary Figure 1A**). Following footprinting, PacBio sequencing, and computational processing, we observed near-complete chromatinization of our DNA templates across the entire length of each footprinted molecule (note paucity of purple ‘accessible’ bases, **Figure 1F**). Methyltransferase-inaccessible footprint sizes corresponded with expected mono- and di-nucleosome footprints (**Figure 1G**), and we estimated a density of 67 ± 3 nucleosomes per molecule (median ± median absolute deviation), confirming full coverage (**Figure 1H**). Strikingly, we observed an enrichment for short (< 80 bp) footprints around *ARS1*, with a corresponding decrease in nucleosomal (80 - 160 bp) and oligonucleosomal (160+ bp) footprints (**Figure 1I**; top), indicative of ORC loading at the replication origin (schematic of origin in **Supplementary Figure 1B**); these patterns were also visible at single-molecule resolution (**Figure 1I**; bottom). Control footprinting experiments on non-chromatinized DNA molecules incubated with replisome components also demonstrated similar footprints aligned at the *ARS1* sites (**Supplementary Figures 1C,D**), confirming that these are dependent on replication proteins and not the result of footprinting short chromatin protections such as H2A-H2B dimers. Together, these analyses demonstrate the quality of our input chromatin preparation, confirm the loading of ORC on *ARS1* to mark origin recognition, and establish SAMOSA-ChAAT’s ability to structurally characterize chromatin organization from these *in vitro* reactions.

### Computational identification of replicated fibers from footprinted reconstitutions

We recently developed <u>R</u>eplication <u>A</u>ware <u>S</u>ingle-molecule <u>A</u>ccessibility <u>M</u>apping (RASAM) (*27*), which measures single-molecule chromatin accessibility and incorporation of BrdUTP on chromatin fibers. We hypothesized that this approach could be adapted to study chromatin replication *in vitro*. To this end, we carried out all reactions with 100% BrdUTP replacing dTTP, thus marking replicated molecules. BrdUTP use did not impact DNA replication through chromatin (**Supplementary Figure 2A**). We then developed a computational pipeline to estimate BrdUTP incorporation on Watson (blue) or Crick (green) strands from single-molecule sequencer kinetics (**Figure 2A**) (*27, 33, 37*). Unsupervised Leiden (*38*) clustering of each molecule separated by strand led to distinct patterns on chromatinized templates (**Figure 2B**): asymmetric clusters reflected replicated molecules containing either Watson or Crick BrdUTP^(+)^ strands, while symmetric clusters reflected molecules that did not incorporate BrdUTP. Importantly, the abundance of asymmetric molecules tracked closely with biochemical estimates of replication efficiency from these reconstitutions (**Supplementary Figure 2B;** 0.77% compared to 1.52% estimated by gel). Finally, this pipeline was robust to multiple different sample types, including footprinted, replicated DNA experiments (**Supplementary Figure 3A**), unmethylated replicated DNA experiments (**Supplementary Figure 3B**), and separate chromatin replication reconstitutions (**Supplementary Figure 3C**).

**Figure 2:**
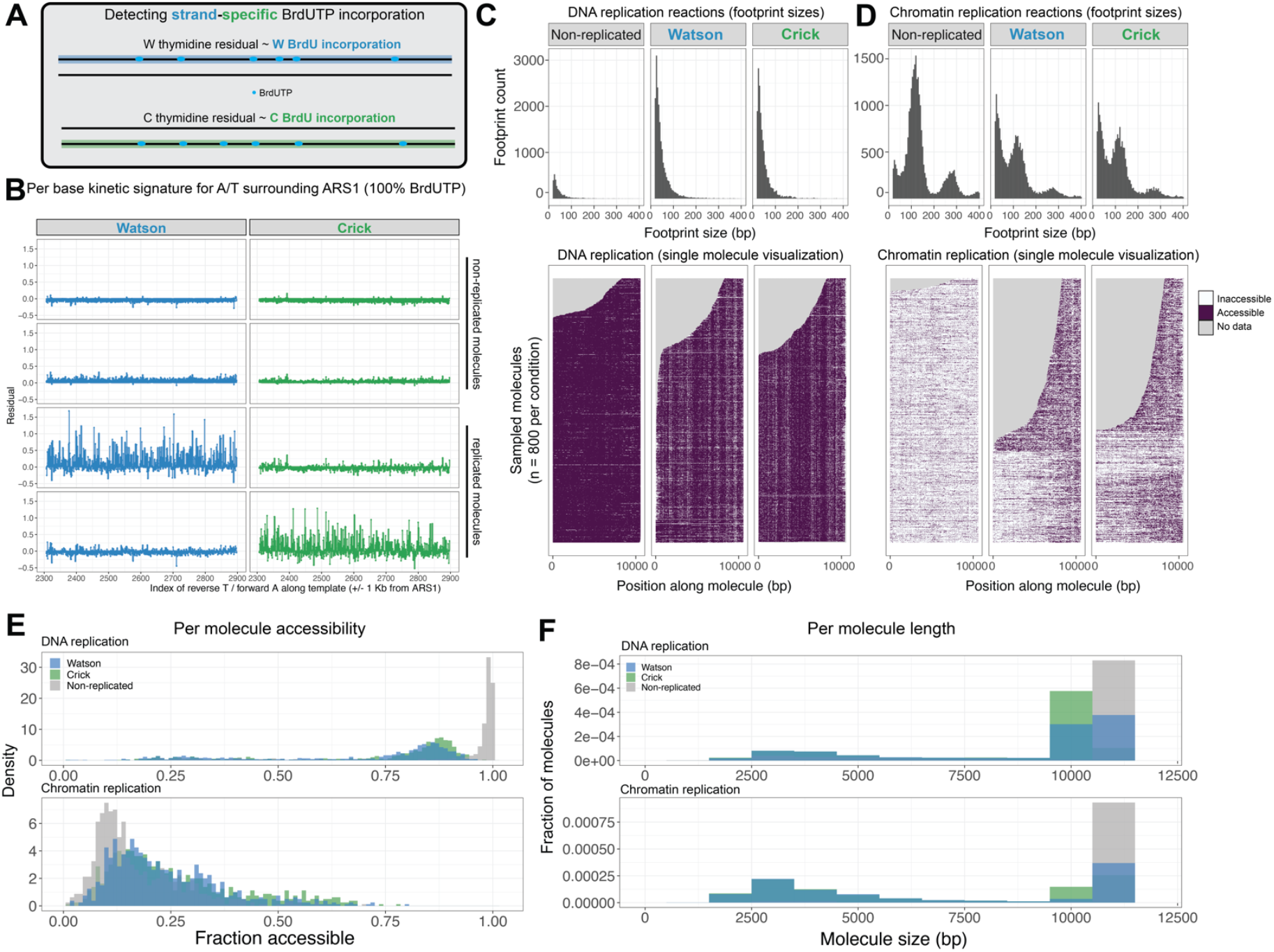
Strand-specific detection of BrdUTP *in silico* and single-molecule characterization of replicated chromatin fibers. **(A)** Schematic of the approach used to detect BrdUTP incorporation on replicated DNA/chromatin with strand-resolution. Alteration in PacBio sequencing polymerase kinetics are reproducibly altered by the presence of nucleotide analogs in sequenced DNA. We apply neural networks to learn the expected kinetics of the sequencing polymerase on unmodified DNA and then compute strand-specific residuals for Watson (W) and Crick (C) strands when sequencing modified DNA. (**B)** Leiden clustering of these residuals for individual sequenced molecules results in grouping of molecules on the basis of strand-specific residual patterns. The top two rows represent non-replicated molecules where the residuals are uniformly low on both strands, while the bottom two rows represent replicated chromatin templates where there is evidence of W BrdUTP incorporation (blue) or C BrdUTP incorporation (green). **(C-D)** Footprint length histograms (top) and single-molecule visualization (bottom) of non-replicated, W BrdUTP, and C BrdUTP molecules from DNA **(C)** or chromatin **(D)** replication reactions. **(E)** Histogram (bin width = 0.01) of per-molecule accessibility for DNA replication (top) and chromatin replication (bottom) reactions for W, C, and non-replicated molecules plotted in **C-D). (F)** Histogram (bin width = 1000 bp) of sequenced molecule sizes (for replicated molecules, ‘replicons’) for W,C, and non-replicated molecules plotted in **C-D)**.

**Figure 3:**
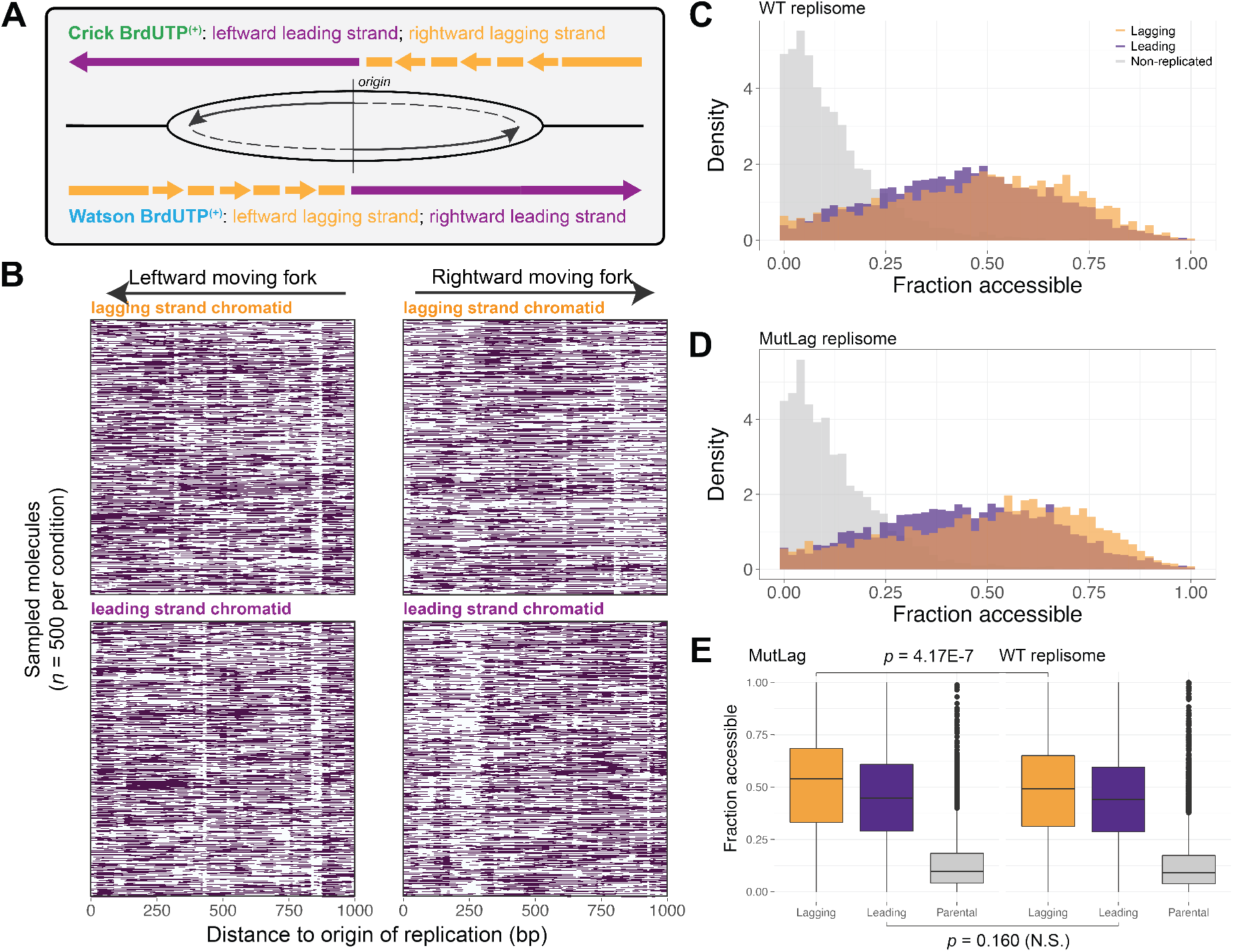
Single-molecule footprinting of replicated lagging- and leading-strand chromatids from reconstituted reactions. **(A)** Schematic of leading and lagging strand assignments, based on the *ARS1* origin position. **(B)** Sampled single-molecule visualization spanning 1 kb positions around the *ARS1* origin at position 0, showing left- and rightward moving forks. BrdUTP^(+)^ molecules obtained from Watson or Crick clusters where the *ARS1* position is known are assigned to either leading or lagging strands **(C)** Accessibility distribution given by the average methylation of single molecules of wildtype replisome reconstitution, separated by non-replicated (grey), lagging strand (orange) and leading strand (purple) classifications.**(D)** Accessibility distribution given by the average methylation of single molecules of mutant lagging-strand (Mcm2–3A and Pol1-2A2) replisome reconstitution, separated by non-replicated (grey), lagging strand (orange) and leading strand (purple) classifications.. **(E)** Boxplot quantifying the mean accessibility from the distributions as plotted in **(C)** and **(D)**, and *p-value* determined by Wilcoxon Rank-Sum test significance level.

We used this pipeline to perform strand-specific footprint size analyses and single-molecule visualization (**Figure 2C,D**). Watson- and Crick-replicated molecules harbored fewer mononucleosome and dinucleosome footprints compared to non-replicated (i.e. parental) molecules, leading to increased single-molecule chromatin accessibility of these fibers following DNA replication (**Figure 2E**). Replicated molecules showed an enrichment for shorter footprints consistent with subnucleosomal particles (*i*.*e*. hexasome; tetrasome), as well as very short protections likely due to replication proteins or altered DNA structures (*e*.*g*. PCNA; G-quadruplexes). We note multiple technical points regarding these data: first, we did not observe a cluster suggestive of re-replication (*i*.*e*. elevated variance on both strands), in line with these reconstitutions promoting a single round of DNA replication (*28*). Second, replicated molecules from chromatin reconstitutions appeared shorter than the ones retrieved from naked DNA replication reactions (**Figure 2F**; gel demonstrating increased replication intermediates **Supplementary Figure 4A**), suggesting that chromatin hinders replisome progression, despite the presence of FACT in our reactions. Third, full-length replicated molecules from chromatin reactions were visually extremely non-uniform (see low accessibility on 5’-end compared to 3’-end on single-molecule visualization for **Figure 2C,D**) and did not have evidence of BrdUTP incorporation on both termini (**Supplementary Figure 4B**). These molecules are thus likely mis-processed products of Pif1/Cdc9/Fen1 (*30*). Together, these data demonstrate our ability to computationally identify and structurally map replicated daughter chromatids from these reconstitutions with strand-specific resolution. These data also highlight the much-added complexity of reconstituting replication of long chromatin templates compared to free DNA or short chromatin templates.

**Figure 4:**
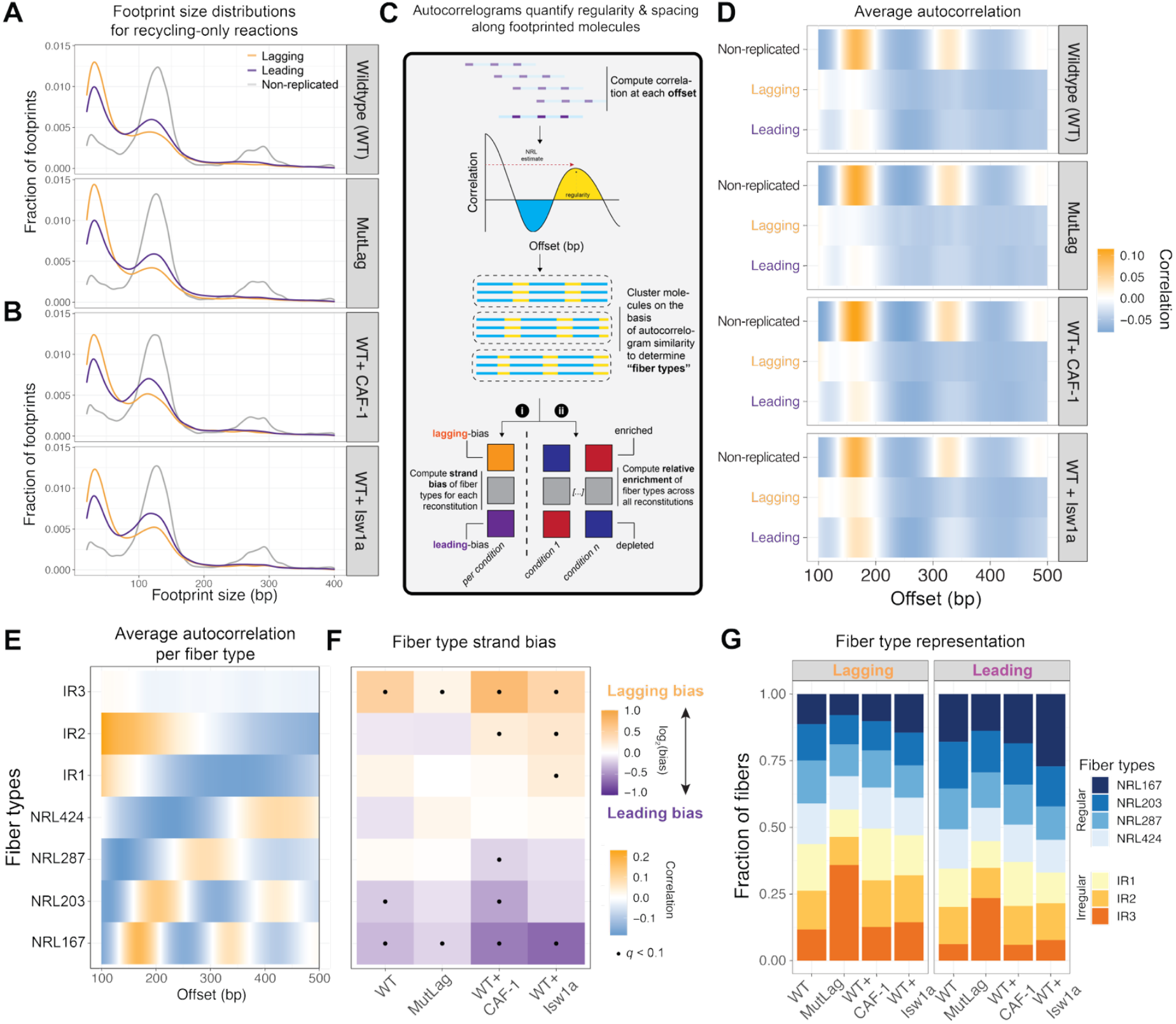
Replisomes recycle histones to generate structurally altered nucleosomes and subnucleosomes with distinct spacing patterns on lagging- and leading-strand chromatids. **(A)** Footprint size distributions plotted as kernel density estimates for wildtype replisome, mutant lagging-strand replisome. **(B)** Same as (A) for wildtype replisome + CAF-1 without donor histone, and wildtype replisome + the ATP-dependent chromatin remodeler Isw1a. **(C)** Visual schematic of the approach used to determine “fiber types” and relative enrichment by quantifying spacing and regularity of chromatin fibers from footprinted molecules. Leiden clustering was used to cluster footprinted molecules based on similarity, to generate chromatin “fiber types” characteristic of different reconstitutions. Once the fiber groups are defined, a series of Fisher’s exact tests are performed, to compute the relative enrichment of chromatin fibers across different reconstitutions (blue-red heatmaps), and strands within each reconstitution (purple-orange heatmaps). **(D)** Average autocorrelogram using Leiden clustering for lagging- and leading-strand chromatin and sampled non-replicated molecules for wildtype replisome, mutant lagging-strand replisome, wildtype replisome + CAF-1 alone and wildtype replisome + ISW1a alone. **(E)** Single-molecule autocorrelations, followed by Leiden clustering across replicated fibers from four reconstitutions: wildtype replisome, mutant lagging-strand replisome, wildtype replisome + CAF-1 alone and wildtype replisome + ISW1a alone, displaying seven fiber types characteristic of these reconstitutions. The fiber types are manually annotated as regular, with specific nucleosome repeat lengths (NRLs) as determined by distance between peaks in signal, or irregular (IR), where either a weak or no autocorrelogram peak was observed. **(F)** Fiber type enrichment determined by Fisher’s exact test, to characterize fiber type representation by strand. **(G)** Relative quantification of fiber type representation per reconstitution for leading and lagging strands.

### Lagging-strand parental histone recycling mechanisms are intrinsic to the replisome

Our footprinting data from purely reconstituted chromatin replication reactions offer a unique opportunity to visualize how the replisome assembles nascent chromatin absent compensatory nuclear factors. As our method identifies BrdUTP incorporation strand and *ARS1* position is known (**Figure 3A**), we next studied how parental histone recycling is carried out by the replisome on individual lagging- and leading-strand chromatids. For all subsequent analyses, we focused our analyses on a 2 kilobase window surrounding *ARS1*. We visualized these molecules separated by both strand- and fork-direction (**Figure 3B**). We also examined the average methylation of molecules from both fork directions, separated by lagging (orange) and leading (purple) classifications, and plotted these values against those for non-replicated chromatin (grey). In line with the sole presence of histone recycling pathways and the omission of *de novo* chromatin assembly proteins, both lagging- and leading-strand chromatids were markedly more accessible compared to non-replicated chromatin fibers (**Figure 3C** and repeat shown in **Supplementary Figure 5A**). Importantly, lagging and leading strands showed comparable opening effects with a slight preference for leading strand assembly (**Figure 3C,E** and repeat shown in **Supplementary Figure 5A-B**), supporting the current model that parental histone recycling almost equally distributes nucleosomes to both replicated strands (*12*). These data represent the first views of protein-DNA interactions on purely reconstituted lagging- and leading-strand chromatids synthesized by the yeast replisome.

**Figure 5:**
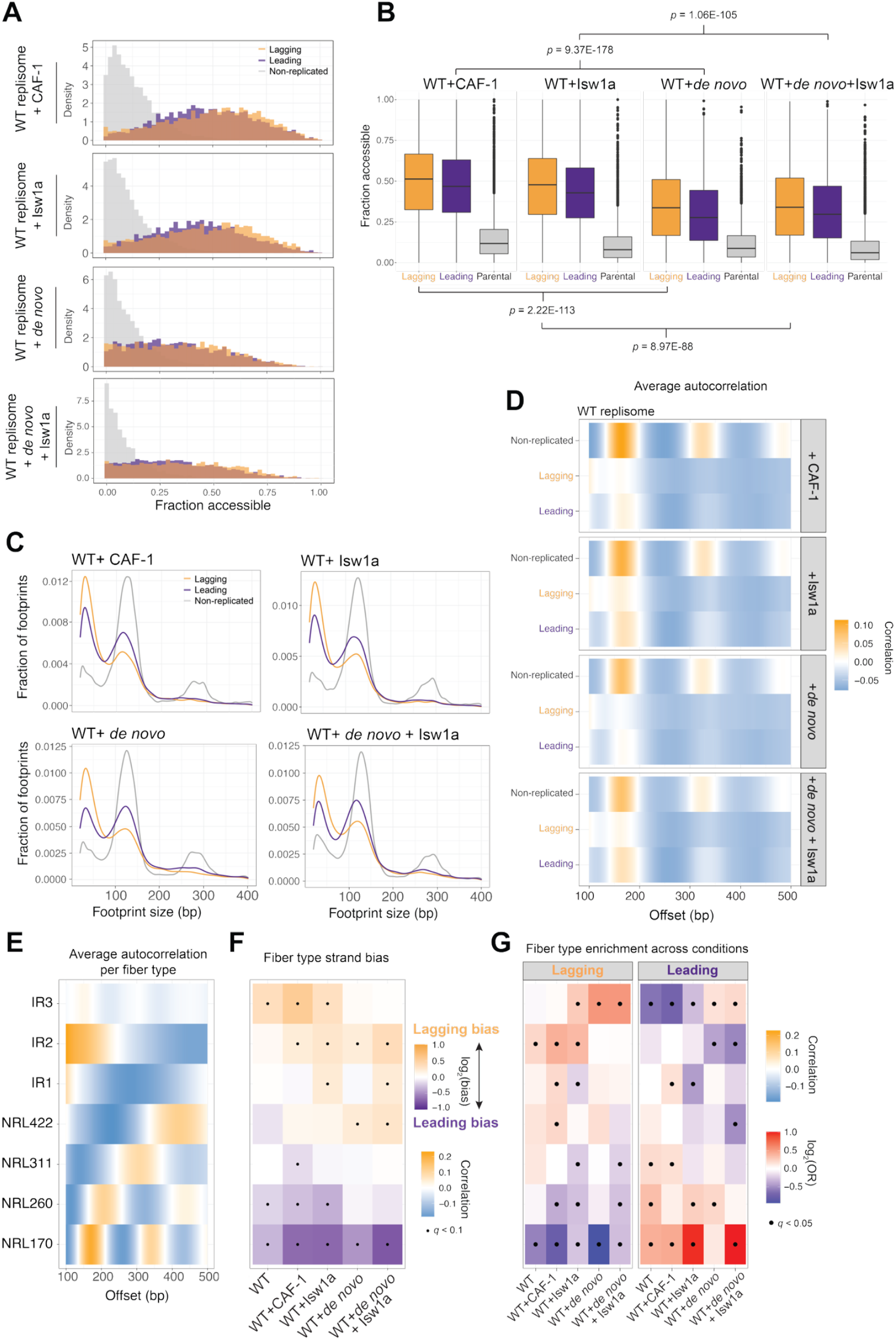
Single-molecule analyses of reconstituted *de novo* deposition and remodeling replication reactions. **(A)** Accessibility distribution given by the average methylation of single molecules, separated by non-replicated (grey), lagging strand (orange) and leading strand (purple) classifications, representing four reconstitutions: wildtype replisome + CAF-1 alone, wildtype replisome + ISW1a alone, wildtype replisome + *de novo*, wildtype replisome + *de novo* + ISW1a. All *de novo* conditions include soluble histones. **(B)** Boxplot quantifying the mean accessibility from the distributions as plotted in **(A)** and *p-value* determined by Wilcoxon Rank-Sum test significance level. **(C)** Footprint size distributions plotted as kernel density estimates for wildtype replisome + CAF-1, wildtype replisome + ISW1a, wildtype replisome + *de novo*, wildtype replisome + *de novo* + ISW1a. **(D)** Average autocorrelogram using Leiden clustering for lagging- and leading-strand chromatin and sampled non-replicated molecules for wildtype replisome + CAF-1, wildtype replisome + ISW1a, wildtype replisome + *de novo*, wildtype replisome + *de novo* + ISW1a. **(E)** Single-molecule autocorrelations, followed by Leiden clustering across replicated fibers from wildtype replisome, wildtype replisome + CAF-1, wildtype replisome + ISW1a, wildtype replisome + *de novo*, wildtype replisome + *de novo* + ISW1a, displaying seven fiber types characteristic of these reconstitutions. The fiber types are manually annotated as regular, with specific nucleosome repeat lengths (NRLs), or irregular (IR), where either a weak / no autocorrelogram peak was observed. **(F)** Fiber type enrichment determined by Fisher’s exact test, to characterize fiber type representation by strand. **(G)** Fiber type enrichment determined by Fisher’s exact test, to characterize fiber type representation across those different reconstitutions for leading-and lagging-strand.

Cellular studies have identified the Mcm2 and PolA proteins as histone chaperones responsible for histone recycling specifically to the lagging strand (*12–15*). To further test whether our reconstitutions could recapitulate chromatin recycling mechanisms observed in yeast and mammalian cells, we performed chromatin replication and footprinting experiments in the presence of histone-binding mutant for Mcm2 (Mcm2–3A: Y79A Y82A Y91A) and PolA (Pol1-2A2: F58A D62A), hereafter, ‘MutLag’ (**Supplementary Figure 5C**,**D)** (*10, 15*). These mutations did not substantially impact DNA replication (**Supplementary Figure 5D**). However, we did detect significantly increased (Wilcoxon Rank-Sum *p* = 4.17E-7) chromatin opening on specifically lagging strands compared to wildtype control (**Figure 3D-E** and repeat shown in **Supplementary Figure 5E**,**F)**, indicating that Mcm2 and PolA histone binding domains contribute to histone deposition specifically on the lagging strand in our reconstitutions, as seen in cells. These data show that our method enables quantification of single-molecule chromatin accessibility in reconstituted reactions with strand-specific resolution.

### The replisome creates distinct chromatin organization on lagging and leading strands

We next assessed how lagging-strand, leading-strand, and non-replicated chromatin differ at the levels of nucleosome wrapping and spacing. We first performed footprint size analyses for lagging and leading versus parental chromatin (**Figure 4A**). Interestingly, these analyses showed distinct patterns between leading and lagging strand molecules. On the lagging strand, we observed increased representation of 20 – 80 bp (*i*.*e*. ‘sub-nucleosomal’) footprints compared to leading strands and non-replicated molecules (**Figure 4A** and repeats shown in **Supplementary Figure 6A**). This difference was not observed in replication reactions run on naked DNA (**Supplementary Figure 6B**), indicating that these footprints are products of chromatin-related mechanisms (*i*.*e*. deposited tetrasomes, hexasomes, and breathable nucleosomes). Addition of the chromatin assembly factor CAF-1 alone in absence of soluble histones (*i*.*e*. WT+CAF-1), or the ATP-dependent chromatin remodeler Isw1a alone without soluble histones (*i*.*e*. WT+Isw1a) did not alter strand-specific differences (**Figure 4B**). These data suggest that chromatin recycling pathways generate distinctly organized lagging- and leading-strand chromatids.

**Figure 6:**
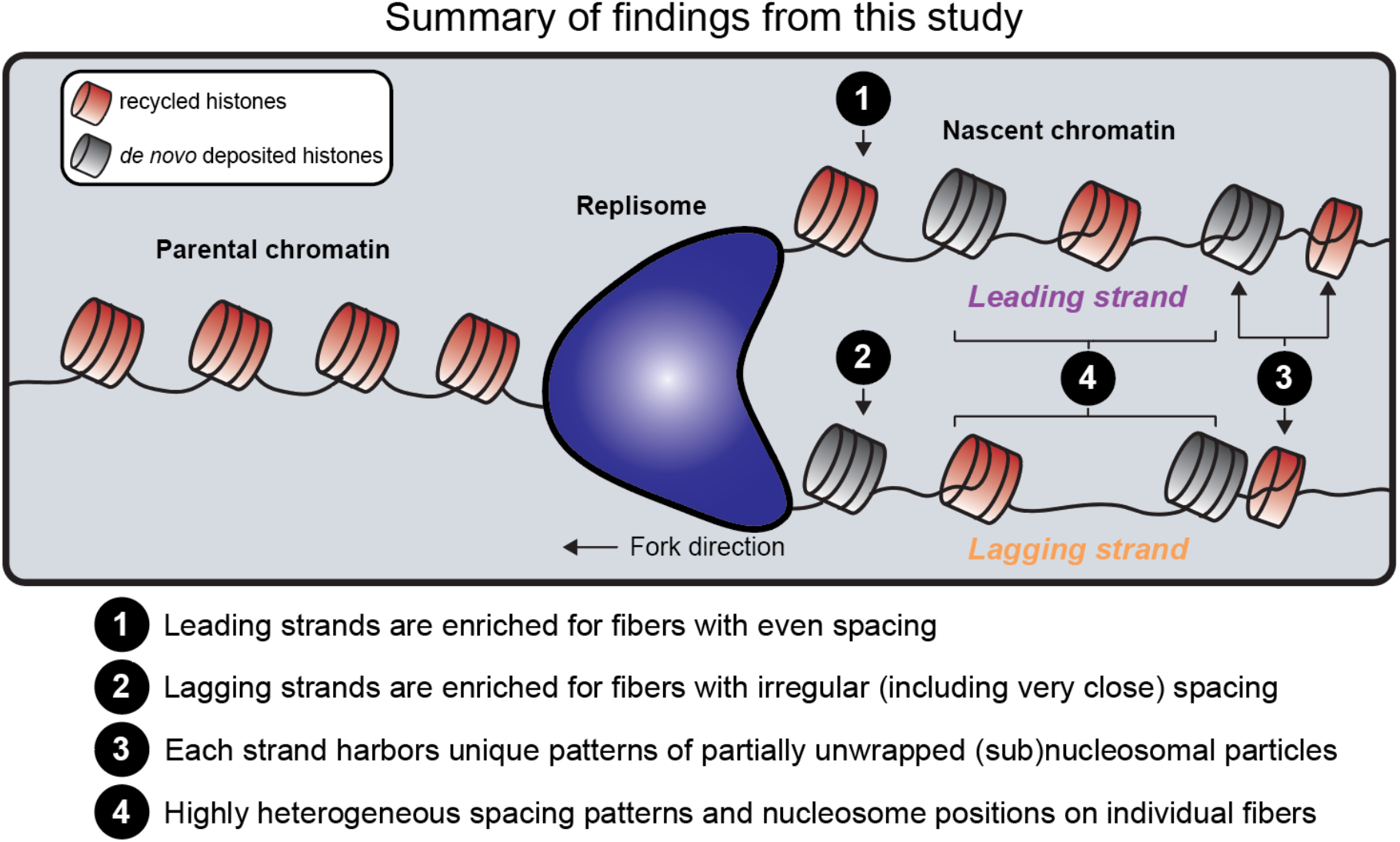
A model for replisome-coupled chromatin fiber assembly based on the findings of this study. From our single-molecule footprinting experiments on reconstituted chromatin fiber replication reactions, we arrive at four main conclusions. **(1)** and **(2):** Leading and lagging strand chromatids are enriched for unique patterns of subnucleosomal particle enrichment and spacing; leading strand chromatin is more likely to harbor evenly spaced particles, while lagging strand chromatin is enriched for more subnucleosomal particles that are irregularly spaced on DNA. Importantly, the addition of the *de novo* deposition pathway and the ATP-dependent chromatin remodeler only partially rescues this “ground state asymmetry,” implying the necessity of more factors for re-establishing symmetric chromatin structure. **(3)** Interestingly, in addition to subnucleosomal particles, we observe partially unwrapped nucleosomes on nascent DNA. **4)** We close by noting that fiber spacing patterns are highly heterogeneous at the single-molecule level: while computed biases represent statistically significant enrichment or depletion of particular structures in our experiments, we also observe *e*.*g*. irregular chromatin fiber types on leading strands and regular chromatin fiber types on lagging strands, implying that the replisome and associated factors are capable of generating these patterns on both strands.

To further investigate differences, we performed single-molecule autocorrelation analysis (**Figure 4C**)(*27, 33, 39*), which assesses nucleosome spacing and regularity on each footprinted molecule. We again used Leiden clustering to define groups of molecules with similar autocorrelation patterns, which we termed ‘fiber types.’ We first examined the average autocorrelation patterns for lagging- and leading-strand chromatin, compared against sampled non-replicated molecules as a control (**Figure 4D**). Non-replicated chromatin arrays were highly regular (see gold ‘heat’ centered at 175 bp) compared to replicated arrays (note less intense and broader offset peaks in heatmaps, **Figure 4D**). Lagging strand chromatin peak positions (which reflect the average spacing between footprints on molecules) were slightly left-shifted compared to leading-strand peak positions (**Figure 4D**), indicating that constituent molecules harbor slightly closer-spaced footprints. Moreover, lagging-strand peak heights were slightly lower compared to leading-strand peak heights (note less-bright gold signal, **Figure 4D**), suggesting a lower abundance of molecules with regularly spaced footprints.

Given the similarity between the average autocorrelation profiles for replicated molecules, we next quantified differences at single-molecule resolution across both strand and reconstitution. We performed Leiden clustering on autocorrelograms for all replicated molecules, resulting in 7 fiber types (**Figure 4E**) that we manually annotated as either: i.) regular with specific nucleosome repeat lengths (NRLs), or ii.) irregular (IR) based on observing either a weak or no autocorrelogram peak. We then performed a series of false discovery rate (Storey’s method(*40*)) corrected Fisher’s exact tests to determine the relative enrichment and depletion of fiber types for each sample between strands (purple-orange heatmaps; *i*.*e*. ‘bias’) and across both samples and strands (blue-red heatmaps; effect sizes presented as odds ratios [O.R.]). We visualized how bias per fiber type differed by strand within samples (**Figure 4F**) and across all samples analyzed together (**Supplementary Figure 6C**). This higher-resolution analysis uncovered clear differences between lagging versus leading strand chromatin structure. Lagging strand molecules were uniformly and significantly biased for irregular clusters such as IR3 (*e*.*g*. WT reconstitution Storey’s *q* = 1.10E-7; log_2_(bias) = 0.152), while leading strand molecules were significantly biased towards more regular fibers such as NRL167 (*e*.*g*. WT reconstitution *q* = 9.94E-7; log_2_(bias) = −0.0953). These patterns also varied when quantified across both strand and sample: WT+Isw1a (in absence of soluble histones), for instance, was more likely to generate NRL167 fibers on leading strands compared to lagging strands (Lagging: *q* = 0.913 [N.S.], O.R. = 1.01; Leading: *q* = 7.41E-49, O.R. = 2.04); this was in contrast to CAF-1 without histones (*i*.*e. ‘*WT+CAF-1’) control samples, which were depleted for NRL167 fibers on lagging strands and enriched for this structure on leading strands (Lagging: *q* = 9.63E-12, O.R. = 0.708; Leading: *q* = 7.04E-11, O.R. = 1.37). Mutations impacting Mcm2 / PolA histone chaperone function (*i*.*e*. ‘MutLag’) did not affect our observation of asymmetry but did lead to significant over-representation of IR3 fibers (Lagging: *q* = 3.71E-165, O.R. = 2.75; Leading: *q =* 7.41E-49, O.R. = 1.69). We stress, however, that these biases and enrichments are computed over a highly heterogeneous mixture of single-molecule states: bar-chart visualization of fiber type representations per-sample (**Figure 4G**) show that all strands and samples sample from each fiber type. Taken together, these data suggest that the yeast replisome can generate ordered and disordered chromatin fiber structures on both lagging- and leading-strands, but that replisome-intrinsic mechanisms are tuned to generate distinct chromatin organization patterns on each strand. Thus, our data reveal an unexpected asymmetric “ground state” for nascent chromatin organization.

### The *de novo* histone deposition pathway and chromatin remodeling partially restore symmetry on lagging- and leading-strand chromatids

In cells, chromatin structure behind the fork is very rapidly modified to generate symmetric lagging and leading strand chromatids. How, then, might this reconstituted ground state be remodeled to restore symmetry? We hypothesized that *de novo* chromatin assembly by histone chaperones CAF-1 and ASF1 and the activities of various chromatin remodeling factors perform this role. To quantify how these pathways remodel daughter chromatids, we carried out experiments adding (1) *de novo* histone deposition factors with soluble histones, and (2) *de novo* histone deposition factors with soluble histones and the ATP-dependent nucleosome remodeler Isw1a. We first examined how per-molecule accessibility is impacted by these additions. Addition of *de novo* histone deposition, with or without Isw1a, visibly and significantly decreased the accessibility of both daughter strands compared to samples where CAF-1 or Isw1a were present without additional histones (**Figure 5A,B**; Wilcoxon Rank-Sum *p* = 9.37E-178 for WT+CAF-1 versus WT+*de novo*; *p* = 1.06E-105 for WT+Isw1a versus WT+*de novo*+Isw1a), supporting enhanced nucleosome assembly on both nascent strands. In reactions performed using MutLag mutants, we observed comparable reduction in chromatin accessibility, though the increased opening of the lagging strand was not completely rescued (**Supplementary Figure 7A-C**). These data confirm that addition of *de novo* histone deposition factors including soluble histones increases chromatinization of molecules in our system (in line with our biochemical data; **Figure 1E**). However, our data also suggest that *de novo* chromatin assembly is not sufficient to completely rescue the impacts of dysregulated lagging strand recycling.

We next assessed the size of nucleosomal footprints for these samples. Footprint analysis showed an enhancement in the mono-nucleosomal peak, particularly when both *de novo* chromatin assembly factors and Isw1a were added to the reaction (**Figure 5C**), suggesting that recombinant *de novo* chromatin assembly factors are sufficient to generate full nucleosomes on nascent DNA generated by the yeast replisome. Interestingly, the addition of *de novo* deposition and Isw1a led to a pronounced leftward shift in the size of nucleosomal protections (particularly on leading strand chromatin), suggestive of systematic nucleosome unwrapping. This shift is evocative of patterns observed by our group on nascent mouse and human chromatin (*27*), possibly coupling ISWI remodeling of nascent nucleosomes in cells (*41*) to nascent nucleosome hyperaccessibility.

Next, we asked how *de novo* histone deposition and Isw1a impact lagging / leading strand fiber structural asymmetry. Average autocorrelation profiles showed more defined nucleosome spacing, particularly in reactions containing Isw1a (**Figure 5D**; see higher intensity gold ‘peaks’ in data). However, leading and lagging strands remained distinctly organized in all experiments (**Figure 5D**), indicating that neither *de novo* chromatin assembly nor Isw1a remodeling can completely neutralize asymmetric chromatin organization. Clustering of single lagging- or leading-strand molecules from these experiments further confirmed these observations, generating distinct fiber types (Figure 5E), with a less pronounced but still significant bias of lagging strand molecules towards more irregular chromatin fibers compared to the more organized leading strand (**Figure 5F**; for example, for WT+CAF-1 samples, Leading: NRL170 *q* = 9.93E-12 & log_2_(bias) = −0.668; Lagging: IR2 *q* = 9.03E-3 & log_2_(bias) = 0.210). Notably, Isw1a-containing reactions with *de novo* histone deposition (‘WT+de novo+Isw1a’) showed increased representation of NRL170 fibers, particularly on the leading strand (**Figure 5F**; NRL170 *q* = 3.80E-12 & log_2_(bias) = −0.763) and also in comparison with other reconstitutions (**Figure 5G**; NRL170 Leading: *q* = 1.00E-36 & O.R. = 2.02). This suggests that ISWI remodelers work differently on lagging and leading strand chromatin, even in the presence of *de novo* deposited nucleosomes and ATP-dependent nucleosome sliding. However, we again stress that these enrichments are computed out of a highly heterogeneous set of fiber types for each sample (**Supplementary Figure 7D**), highlighting the fact that *de novo* deposition and remodeling can generate the full range of chromatin fiber types on both nascent strands. Taken as a whole, our data demonstrate that *de novo* deposition and remodeling promote the formation (and potentially, maturation) of nucleosomes in these reconstitutions, but that these processes are not sufficient to generate symmetric organization of lagging and leading strand chromatin.

## DISCUSSION

### Dissecting chromatin replication *in vitro* through single-molecule footprinting

Here, we lay groundwork for the high-throughput, single-molecule biochemical dissection of chromatin replication reactions. By combining the SAMOSA-ChAAT assay with the highly complex reconstitution of chromatin replication by purified *S. cerevisiae* proteins, we provide the first single-molecule views of histone recycling and *de novo* deposition on kilobase-scale reconstituted chromatin fibers. Our work sets the stage for quantitative dissection of replication initiation, elongation, and termination on chromatinized templates of any DNA sequence of interest, and study of the effects of *trans*-acting factors including different histone chaperones, regulatory factors, and mutant proteins. Importantly, the resolution obtained in this study is considerably higher than what has been possible in cellular systems, allowing for strand-resolved views of chromatin fiber structure in precisely defined conditions uncoupled from gene essentiality.

### Asymmetric organization of nascent chromatin fibers

DNA replication of lagging and leading strands is inherently asymmetric. Our work shows that chromatin generated by the replisome during DNA replication is also inherently asymmetric (**Figure 6**). Lagging and leading strands are differentially organized at the chromatin level: lagging strands are significantly enriched for more irregular nucleosome spacings, and poorer-defined mono-nucleosomal protections; leading strands are significantly enriched for more regular nucleosome spacings, and more precisely defined mono-nucleosomal protections. Our data thus extend observations from over a decade ago on the intrinsic coupling of lagging strand synthesis with chromatin assembly (*42*), by showing that the fiber structures generated by yeast replisomes alone are inherently different. Still, we note that our experiments demonstrate substantial heterogeneity: the yeast replisome is capable of generating a wide variety of ordered and disordered structural patterns on both strands, and the relative distributions of these patterns is shifted by lagging-versus leading-strand mechanisms. Recent studies have established distinct histone recycling mechanisms responsible for histone chaperoning at the replisome between the two daughter strands (*10, 12–16, 21–23*). Our study demonstrates that these different recycling pathways also yield distinct chromatin fiber products. Thus, our method provides a much-needed tool for studying how distinct chaperones generate and resolve strand asymmetry. Increasing the complexity of our reconstitutions (*e*.*g*. through addition of more remodelers (*43*), or post-translational modifications such as H3K56ac (*41*)) to more closely model *in vivo* chromatin replication and maturation remains the essential next step.

### Implications for post-replication mechanisms acting on chromatin

What are the implications of the asymmetric ground state characterized here? It is possible that this asymmetry is purely a consequence of strand-specific DNA replication mechanisms, where for example the discontinuity of the lagging strand may foster more stochastic (*i*.*e*. irregular) nucleosome assembly events, compared to the continuous (and, as shown here, more regular) leading strand-coupled assembly. Interestingly, very recent work has identified asymmetry in sister chromatid cohesion (*44*) and in loading of RNA Polymerase II (*45*), highlighting that strand-specific differences in chromatin organization may have functional implications. Speculatively, strand-specific differences in chromatin organization—and their resolution dynamics—could also have functional consequences on replication stress resolution, DNA damage response or even the persistence of mutational lesions over multiple cell divisions (*46–49*). The physiological significance of modeling and resolving asymmetry at a biochemical level highlights the value of the platform we present here: by carrying out chromatin replication in precisely reconstituted reactions, our system enables single-molecule mechanistic dissection of replication-associated phenomena. Ultimately combining our methods with high-resolution profiling methods *in vivo* will elucidate the physiological significance of asymmetric ground states across DNA-templated processes.

## ACKNOWLEDGEMENTS

We thank members of the Mattiroli and Ramani lab for useful discussions. We thank Juan Garaycoechea (Hubrecht Institute) and Hiten Madhani (UCSF) for critical feedback on the manuscript.

## Funding

FM acknowledges funding from the EU (ERC StG 851564), the Dutch Research Organization (NWO VI.Vidi.233.038) and EMBO (yip 5526-2024). VR acknowledges funding from National Institutes of Health grant DP2-HG012442, and through the generous support of the Searle Scholars Program and W.M. Keck Foundation.

## Author Contributions

B.V.E., H.J.R., and I.R. performed experiments, B.V.E., H.J.R., K.R., and V.R. performed analyses, B.E., H.R., V.R., and F.M wrote the original manuscript, all authors contributed to review & editing.

## Competing Interests

The authors declare no competing interests.

## Data and materials availability

All material generated in this study are available on request. Sequencing data have been deposited at the Gene Expression Omnibus under GSEXXXXXX. Custom code used in this study is available in a permanent public repository on GitHub at https://github.com/RamaniLab/repli-SAMOSA-ChAAT.

## SUPPLEMENTARY MATERIALS

Materials and Methods

Figs. S1 to S4

References (*48-52*)

## Notes

### Competing Interest Statement

The authors have declared no competing interest.

## REFERENCES

1. K. R. Stewart-Morgan, N. Petryk, A. Groth, Chromatin replication and epigenetic cell memory. Nat. Cell Biol. 22, 361–371 (2020).

2. O. Willhoft, A. Costa, A structural framework for DNA replication and transcription through chromatin. Curr. Opin. Struct. Biol. 71, 51–58 (2021).

3. W. Du, G. Shi, C.-M. Shan, Z. Li, B. Zhu, S. Jia, Q. Li, Z. Zhang, Mechanisms of chromatin-based epigenetic inheritance. Sci. China Life Sci. 65, 2162–2190 (2022).

4. Z. Li, Z. Zhang, A tale of two strands: Decoding chromatin replication through strand-specific sequencing. Mol. Cell 85, 238–261 (2025).

5. S. Bell, K. Labib, Chromosome Duplication in Saccharomyces cerevisiae. Genetics 203, 1027–1067 (2016).

6. S. Smith, B. Stillman, Purification and characterization of CAF-I, a human cell factor required for chromatin assembly during DNA replication in vitro. Cell 58, 15–25 (1989).

7. G. Almouzni, M. Méchali, A. P. Wolffe, Competition between transcription complex assembly and chromatin assembly on replicating DNA. EMBO J. 9, 573–582 (1990).

8. J. K. Tyler, C. R. Adams, S. R. Chen, R. Kobayashi, R. T. Kamakaka, J. T. Kadonaga, The RCAF complex mediates chromatin assembly during DNA replication and repair. Nature 402, 555–560 (1999).

9. M. Xu, C. Long, X. Chen, C. Huang, S. Chen, B. Zhu, Partitioning of histone H3-H4 tetramers during DNA replication-dependent chromatin assembly. Science 328, 94–98 (2010).

10. M. Foltman, C. Evrin, G. De Piccoli, R. C. Jones, R. D. Edmondson, Y. Katou, R. Nakato, K. Shirahige, K. Labib, Eukaryotic replisome components cooperate to process histones during chromosome replication. Cell Rep. 3, 892–904 (2013).

11. S. Ramachandran, K. Ahmad, S. Henikoff, Capitalizing on disaster: Establishing chromatin specificity behind the replication fork. Bioessays 39 (2017).

12. N. Petryk, M. Dalby, A. Wenger, C. B. Stromme, A. Strandsby, R. Andersson, A. Groth, MCM2 promotes symmetric inheritance of modified histones during DNA replication. Science 361, 1389–1392 (2018).

13. C. Yu, H. Gan, A. Serra-Cardona, L. Zhang, S. Gan, S. Sharma, E. Johansson, A. Chabes, R.-M. Xu, Z. Zhang, A mechanism for preventing asymmetric histone segregation onto replicating DNA strands. Science 361, 1386–1389 (2018).

14. H. Gan, A. Serra-Cardona, X. Hua, H. Zhou, K. Labib, C. Yu, Z. Zhang, The Mcm2-Ctf4-Polα axis facilitates parental histone H3-H4 transfer to lagging strands. Mol. Cell 72, 140-151.e3 (2018).

15. C. Evrin, J. D. Maman, A. Diamante, L. Pellegrini, K. Labib, Histone H2A-H2B binding by Pol α in the eukaryotic replisome contributes to the maintenance of repressive chromatin. EMBO J. 37 (2018).

16. R. Bellelli, O. Belan, V. E. Pye, C. Clement, S. L. Maslen, J. M. Skehel, P. Cherepanov, G. Almouzni, S. J. Boulton, POLE3-POLE4 Is a Histone H3-H4 Chaperone that Maintains Chromatin Integrity during DNA Replication. Mol. Cell 72, 112-126.e5 (2018).

17. B. Safaric, E. Chacin, M. J. Scherr, L. Rajappa, C. Gebhardt, C. F. Kurat, T. Cordes, K. E. Duderstadt, The fork protection complex recruits FACT to reorganize nucleosomes during replication. Nucleic Acids Res. 50, 1317–1334 (2022).

18. C. Rouillon, B. V. Eckhardt, L. Kollenstart, F. Gruss, A. E. E. Verkennis, I. Rondeel, P. H. L. Krijger, G. Ricci, A. Biran, T. van Laar, C. M. Delvaux de Fenffe, G. Luppens, P. Albanese, K. Sato, R. A. Scheltema, W. de Laat, P. Knipscheer, N. H. Dekker, A. Groth, F. Mattiroli, CAF-1 deposits newly synthesized histones during DNA replication using distinct mechanisms on the leading and lagging strands. Nucleic Acids Res. 51, 3770–3792 (2023).

19. C.-P. Liu, Z. Yu, J. Xiong, J. Hu, A. Song, D. Ding, C. Yu, N. Yang, M. Wang, J. Yu, P. Hou, K. Zeng, Z. Li, Z. Zhang, X. Zhang, W. Li, Z. Zhang, B. Zhu, G. Li, R.-M. Xu, Structural insights into histone binding and nucleosome assembly by chromatin assembly factor-1. Science 381, eadd8673 (2023).

20. Z. Li, S. Duan, X. Hua, X. Xu, Y. Li, D. Menolfi, H. Zhou, C. Lu, S. Zha, S. P. Goff, Z. Zhang, Asymmetric distribution of parental H3K9me3 in S phase silences L1 elements. Nature 623, 643–651 (2023).

21. J. Yu, Y. Zhang, Y. Fang, J. A. Paulo, D. Yaghoubi, X. Hua, G. Shipkovenska, T. Toda, Z. Zhang, S. P. Gygi, S. Jia, Q. Li, D. Moazed, A replisome-associated histone H3-H4 chaperone required for epigenetic inheritance. Cell, doi: 10.1016/j.cell.2024.07.006 (2024).

22. S. J. Charlton, V. Flury, Y. Kanoh, A. V. Genzor, L. Kollenstart, W. Ao, P. Brøgger, M. B. Weisser, M. Adamus, N. Alcaraz, C. M. Delvaux de Fenffe, F. Mattiroli, G. Montoya, H. Masai, A. Groth, G. Thon, The fork protection complex promotes parental histone recycling and epigenetic memory. Cell, doi: 10.1016/j.cell.2024.07.017 (2024).

23. T. Toda, Y. Fang, C.-M. Shan, X. Hua, J.-K. Kim, L. C. Tang, M. Jovanovic, L. Tong, F. Qiao, Z. Zhang, S. Jia, Mrc1 regulates parental histone segregation and heterochromatin inheritance. Mol. Cell, doi: 10.1016/j.molcel.2024.07.002 (2024).

24. N. Li, Y. Gao, Y. Zhang, D. Yu, J. Lin, J. Feng, J. Li, Z. Xu, Y. Zhang, S. Dang, K. Zhou, Y. Liu, X. D. Li, B. K. Tye, Q. Li, N. Gao, Y. Zhai, Parental histone transfer caught at the replication fork. Nature 627, 890–897 (2024).

25. J. Dreyer, G. Ricci, J. van den Berg, V. Bhardwaj, J. Funk, C. Armstrong, V. van Batenburg, C. Sine, M. VanInsberghe, R. B. Tjeerdsma, R. Marsman, I. K. Mandemaker, S. di Sanzo, J. Costantini, S. G. Manzo, A. Biran, C. Burny, M. A. T. M. van Vugt, M. Völker-Albert, A. Groth, S. L. Spencer, A. van Oudenaarden, F. Mattiroli, Acute multi-level response to defective de novo chromatin assembly in S-phase. Mol. Cell 84, 4945 (2024).

26. V. Flury, N. Reverón-Gómez, N. Alcaraz, K. R. Stewart-Morgan, A. Wenger, R. J. Klose, A. Groth, Recycling of modified H2A-H2B provides short-term memory of chromatin states. Cell 186, 1050-1065.e19 (2023).

27. M. S. Ostrowski, M. G. Yang, C. P. McNally, N. J. Abdulhay, S. Wang, K. Renduchintala, I. Irkliyenko, Biran, B. T. L. Chew, A. D. Midha, E. V. Wong, J. Sandoval, I. H. Jain, A. Groth, E. P. Nora, H. Goodarzi, V. Ramani, The single-molecule accessibility landscape of newly replicated mammalian chromatin. Cell, doi: 10.1016/j.cell.2024.10.039 (2024).

28. J. T. P. Yeeles, T. D. Deegan, A. Janska, A. Early, J. F. X. Diffley, Regulated eukaryotic DNA replication origin firing with purified proteins. Nature 519, 431–435 (2015).

29. J. T. P. Yeeles, A. Janska, A. Early, J. F. X. Diffley, How the eukaryotic replisome achieves rapid and efficient DNA replication. Mol. Cell 65, 105–116 (2017).

30. T. D. Deegan, J. Baxter, M.Á. Ortiz Bazán, J. T. P. Yeeles, K. P. M. Labib, Pif1-family helicases support fork convergence during DNA replication termination in eukaryotes. Mol. Cell 74, 231-244.e9 (2019).

31. C. F. Kurat, J. T. P. Yeeles, H. Patel, A. Early, J. F. X. Diffley, Chromatin controls DNA replication origin selection, lagging-strand synthesis, and replication fork rates. Mol. Cell 65, 117–130 (2017).

32. S. Devbhandari, J. Jiang, C. Kumar, I. Whitehouse, D. Remus, Chromatin constrains the initiation and elongation of DNA replication. Mol. Cell 65, 131–141 (2017).

33. N. J. Abdulhay, L. J. Hsieh, C. P. McNally, M. S. Ostrowski, C. M. Moore, M. Ketavarapu, S. Kasinathan, S. Nanda, K. Wu, U. S. Chio, Z. Zhou, H. Goodarzi, G. J. Narlikar, V. Ramani, Nucleosome density shapes kilobase-scale regulation by a mammalian chromatin remodeler. Nat. Struct. Mol. Biol., doi: 10.1038/s41594-023-01093-6 (2023).

34. C. Moore, E. Wong, U. Kaur, U. S. Chio, Z. Zhou, M. Ostrowski, K. Wu, I. Irkliyenko, S. Wang, V. Ramani, G. J. Narlikar, ATP-dependent remodeling of chromatin condensates uncovers distinct mesoscale effects of two remodelers, bioRxiv (2024). 10.1101/2024.09.10.611504.

35. D. V. Fyodorov, J. T. Kadonaga, Chromatin assembly in vitro with purified recombinant ACF and NAP-1. Methods Enzymol. 371, 499–515 (2003).

36. I. A. Murray, R. D. Morgan, Y. Luyten, A. Fomenkov, I.R. Corrêa Jr, N. Dai, M. B. Allaw, X. Zhang, X. Cheng, R. J. Roberts, The non-specific adenine DNA methyltransferase M.EcoGII. Nucleic Acids Res. 46, 840–848 (2018).

37. B. A. Flusberg, D. R. Webster, J. H. Lee, K. J. Travers, E. C. Olivares, T. A. Clark, J. Korlach, S. W. Turner, Direct detection of DNA methylation during single-molecule, real-time sequencing. Nat. Methods 7, 461–465 (2010).

38. V. A. Traag, L. Waltman, N. J. van Eck, From Louvain to Leiden: guaranteeing well-connected communities. Sci. Rep. 9, 5233 (2019).

39. N. J. Abdulhay, C. P. McNally, L. J. Hsieh, S. Kasinathan, A. Keith, L. S. Estes, M. Karimzadeh, J. G. Underwood, H. Goodarzi, G. J. Narlikar, V. Ramani, Massively multiplex single-molecule oligonucleosome footprinting. Elife 9 (2020).

40. J. D. Storey, R. Tibshirani, Statistical significance for genomewide studies. Proc. Natl. Acad. Sci. U. S. A. 100, 9440–9445 (2003).

41. S. Duan, I. M. Nodelman, H. Zhou, T. Tsukiyama, G. D. Bowman, Z. Zhang, H3K56 acetylation regulates chromatin maturation following DNA replication. Nat. Commun. 16, 134 (2025).

42. D. J. Smith, I. Whitehouse, Intrinsic coupling of lagging-strand synthesis to chromatin assembly. Nature 483, 434–438 (2012).

43. E. Chacin, K.-U. Reusswig, J. Furtmeier, P. Bansal, L. A. Karl, B. Pfander, T. Straub, P. Korber, C. F. Kurat, Establishment and function of chromatin organization at replication origins. Nature 616, 836–842 (2023).

44. F. Corsi, S. Kolesnikova, T. L. Steinacker, Z. Takács, P. Batty, M. Mitter, D. W. Gerlich, A. Goloborodko, Conformational asymmetry of replicated human chromosomes, bioRxiv (2025). 10.1101/2025.07.09.663929.

45. R. Ziane, A. Camasses, M. Radman-Livaja, The asymmetric distribution of RNA polymerase II and nucleosomes on replicated daughter genomes is caused by differences in replication timing between the lagging and the leading strand. Genome Res. 32, 337–356 (2022).

46. M. A. M. Reijns, H. Kemp, J. Ding, S.M. de Procé, A. Jackson, M. S. Taylor, Lagging strand replication shapes the mutational landscape of the genome. Nature 518, 502–506 (2015).

47. S. J. Aitken, C. Anderson, F. Connor, O. Pich, V. Sundaram, C. Feig, T. F. Rayner, M. Lukk, S. Aitken, J. Luft, E. Kentepozidou, C. Arnedo-Pac, S. Beentjes, S. Davies, R. M. Drews, A. Ewing, V. B. Kaiser, A. Khamseh, E. López-Arribillaga, A. Redmond, J. Santoyo-Lopez, I. Sentís, L. Talmane, A. D. Yates, Sarah J. Stuart Craig J. Claudia Frances Ruben M. Ailith Aitken Aitken Anderson Arnedo-Pac Connor Drews Ewi, S. J. Aitken, S. Aitken, C. Arnedo-Pac, F. Connor, P. Flicek, N. López-Bigas, D. T. Odom, O. Pich, C. A. Semple, M. S. Taylor, C. A. Semple, P. Flicek, D. T. Odom, M. S. Taylor, Pervasive lesion segregation shapes cancer genome evolution. Nature 583, 265–270 (2019).

48. C. González-Garrido, F. Prado, Parental histone distribution and location of the replication obstacle at nascent strands control homologous recombination. Cell Rep. 42, 112174 (2023).

49. M. Spencer Chapman, E. Mitchell, K. Yoshida, N. Williams, M. A. Fabre, A. M. Ranzoni, P. S. Robinson, L. D. Kregar, M. Wilk, S. Boettcher, K. Mahbubani, K. Saeb Parsy, K. H. C. Gowers, S. M. Janes, S. W. K. Ng, M. Hoare, A. R. Green, G. S. Vassiliou, A. Cvejic, M. G. Manz, E. Laurenti, I. Martincorena, M. R. Stratton, J. Nangalia, T. H. H. Coorens, P. J. Campbell, Prolonged persistence of mutagenic DNA lesions in somatic cells. Nature 638, 729–738 (2025).

48. D. Baretić, M. Jenkyn-Bedford, V. Aria, G. Cannone, M. Skehel, J. T. P. Yeeles, Cryo-EM structure of the fork protection complex bound to CMG at a replication fork. Mol. Cell 78, 926-940.e13 (2020).

49. F. Mattiroli, Y. Gu, T. Yadav, J. L. Balsbaugh, M. R. Harris, E. S. Findlay, Y. Liu, C. A. Radebaugh, L. A. Stargell, N. G. Ahn, I. Whitehouse, K. Luger, DNA-mediated association of two histone-bound complexes of yeast Chromatin Assembly Factor-1 (CAF-1) drives tetrasome assembly in the wake of DNA replication. Elife 6 (2017).

50. P. N. Dyer, R. S. Edayathumangalam, C. L. White, Y. Bao, S. Chakravarthy, U. M. Muthurajan, K. Luger, Reconstitution of nucleosome core particles from recombinant histones and DNA. Methods Enzymol. 375, 23–44 (2004).

51. J. C. Vary Jr, T. G. Fazzio, T. Tsukiyama, Assembly of yeast chromatin using ISWI complexes. Methods Enzymol. 375, 88–102 (2004).

52. S. Nanda, K. Wu, I. Irkliyenko, B. Woo, M. S. Ostrowski, A. S. Clugston, L. C. Sayles, L. Xu, A. T. Satpathy, H. G. Nguyen, E. Alejandro Sweet-Cordero, H. Goodarzi, S. Kasinathan, V. Ramani, Direct transposition of native DNA for sensitive multimodal single-molecule sequencing. Nat. Genet., doi: 10.1038/s41588-024-01748-0 (2024).

