## SUPPLEMENTARY MATERIALS for "The eukaryotic replisome intrinsically generates asymmetric daughter chromatin fibers"

This file includes:

Supplementary Figures S1-S7

Materials and Methods

### SUPPLEMENTARY FIGURES

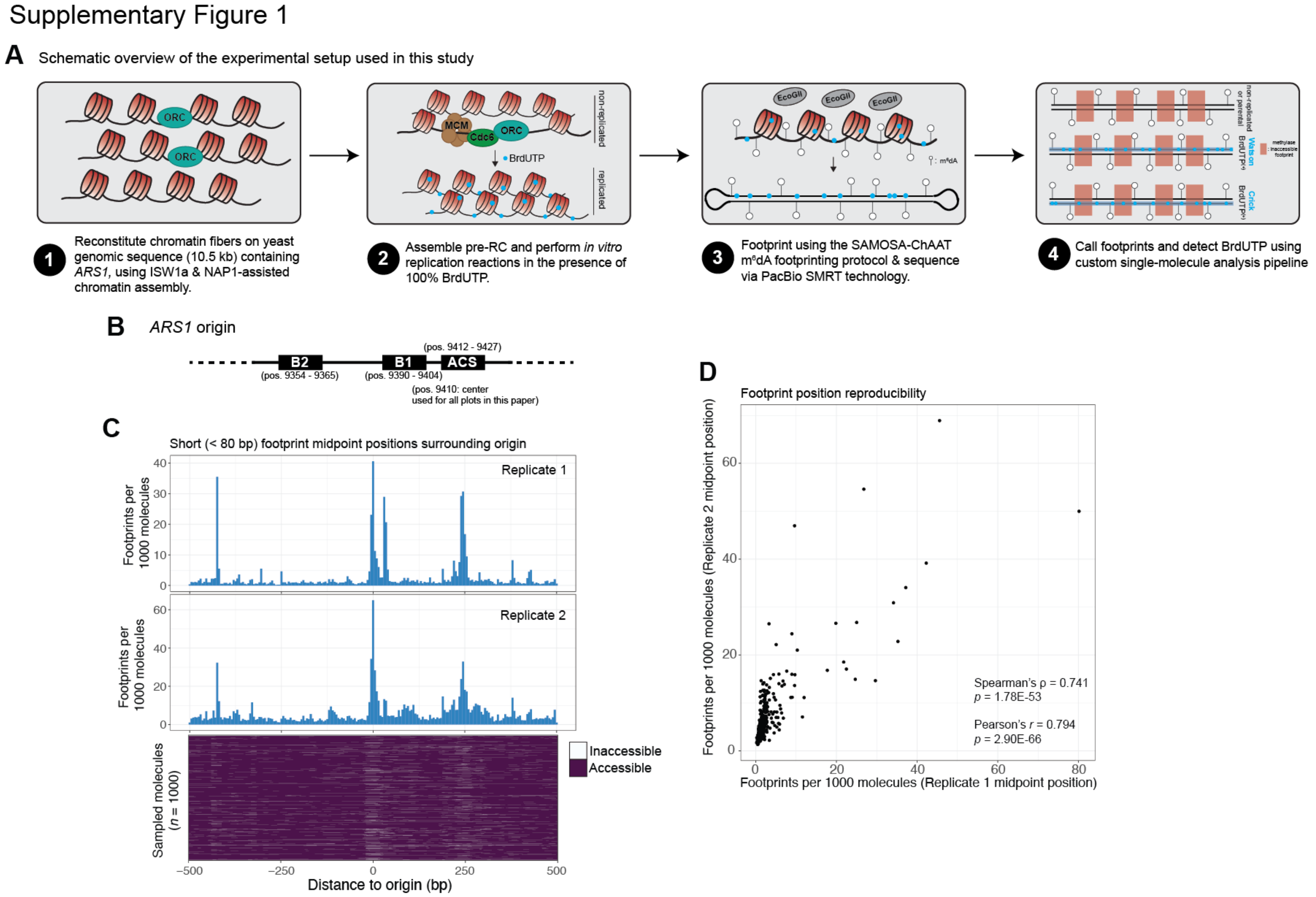


**Supplementary Figure 1: Schematic overview of the SAMOSA-ChAAT pipeline to study chromatin fiber replication and control replication experiments on naked DNA. (A)** Visual schematic of the SAMOSA to test Chromatin Accessibility of Assembled Templates (SAMOSA-ChAAT) approach applied to reconstituted chromatin replication reactions. We first reconstitute DNA replication on chromatinized templates, following the protocol of Kurat *et al*. We initiate replication in the presence of 100% BrdUTP substituted for dTTP and allow replication to occur, resulting in a mixed population of non-replicated fibers and replicated fibers containing BrdUTP. We then perform single-molecule footprinting with the m$^6$dAase EcoGII and sequence resulting molecules using the PacBio single-molecule real-time sequencer. We then use custom computational pipelines to infer the position of methylase-inaccessible footprints and whether BrdUTP has been incorporated on sequenced single molecules. **(B)** Schematic of the *ARS1* region on assembled templates, with the positions (in nucleotides) of the B2, B1, and ACS elements annotated. **(C)** Histogram of footprint midpoint positions (top) and sampled molecules (bottom), oriented with the *ARS1* origin centered, for both replicates. **(D)** Footprint midpoint positions are highly quantitatively reproducible across replicate reconstitution experiments.

Supplementary Figure 2

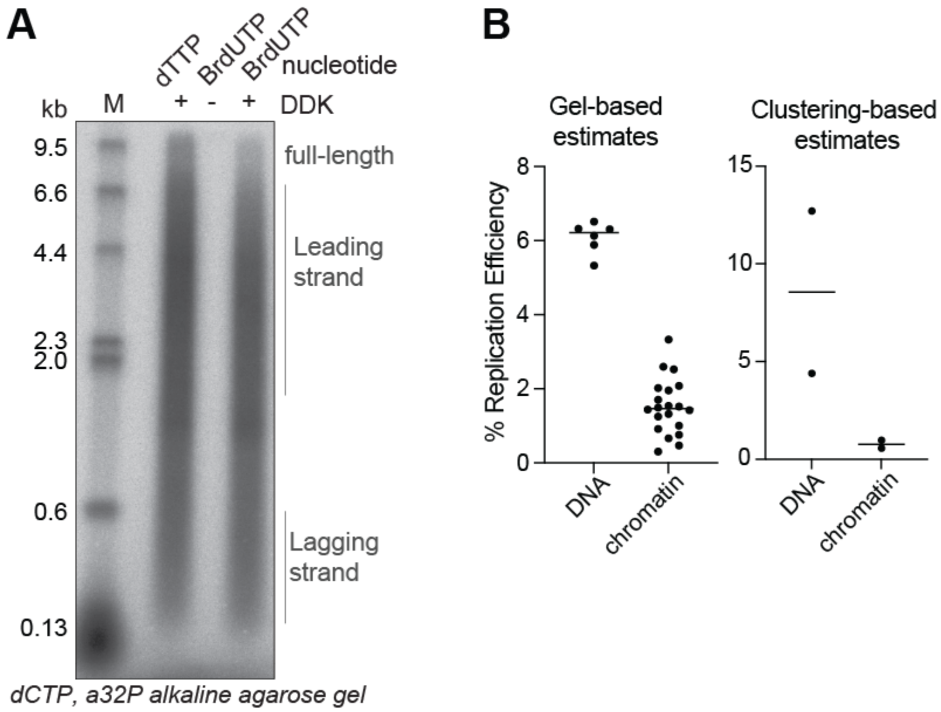


**Supplementary Figure 2: Validation of BrdUTP incorporation and replication efficiencies. (A)** Autoradiography of denaturing agarose gel of reactions performed in presence of either dTTP or BrdUTP. DDK omission is used as control to confirm origin-based replication mechanisms. Replication reactions carried out in the presence of 100% BrdUTP instead of dTTP do not show differences in replication efficiencies. **(B)** quantification of replication efficiency of reactions run on naked DNA or chromatin templates. Left graph shows replication efficiency calculated from biochemical readouts (based on radioactivity incorporation, see methods). Right graph shows the total percentage of replicated molecules found by clustering of PacBio polymerase kinetics to detect BrdUTP incorporation.

Supplementary Figure 3

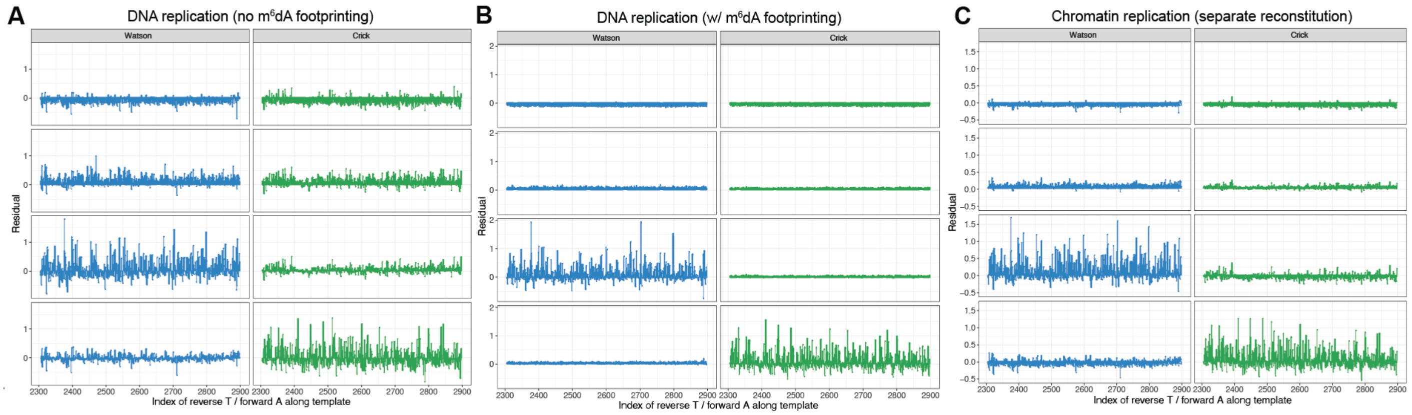


**Supplementary Figure 3: Single-molecule clustering per strand shows asymmetric patterns on Watson or Crick BrdUTP$^{(+)}$ strands in replicated samples.** Unsupervised Leiden clustering of the residuals for individual sequenced molecules results in grouping of molecules on the basis of strand-specific residual patterns. The top two rows include non-replicated molecules where the residuals are uniformly low on both strands, while the bottom two rows represent replicated chromatin templates where there is evidence of W BrdUTP incorporation (blue) or C BrdUTP incorporation (green) for the following reconstitutions: **(A)** Naked DNA replication reaction not footprinted by the m$^6$dA methyltransferase EcoGII **(B)** Naked DNA replication reactions footprinted by the m$^6$dA methyltransferase EcoGII **(C)** Chromatin replication reaction.

### Supplementary Figure 4

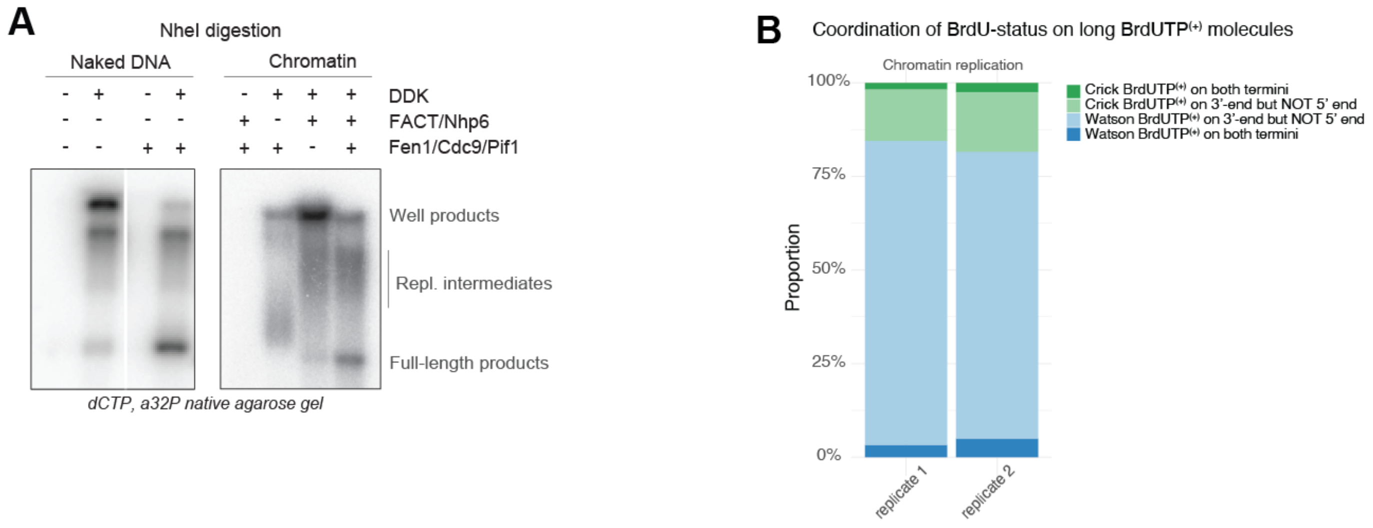


**Supplementary Figure 4: Replicated chromatin fibers are shorter than full-length products suggesting replisome may stall during replication of long chromatin templates. (A)** Autoradiography of native agarose gel of naked DNA or chromatin reaction to show heterogeneity in product formation after termination reaction (note presence of replication intermediates and well products). Chromatin reaction shows decreased efficiency in successful full-length product formation, compared to the naked DNA samples. **(B)** Clustering for BrdUTP signal near the 5' end (distal to *ARS1*) shows that majority of long or near-full length replicated fibers have BrdUTP signal only on the 3' at *ARS1* and are likely ligation products, not long replicated fibers, across two replicates of chromatin replication with only parental histone recycling.

Supplementary Figure 5

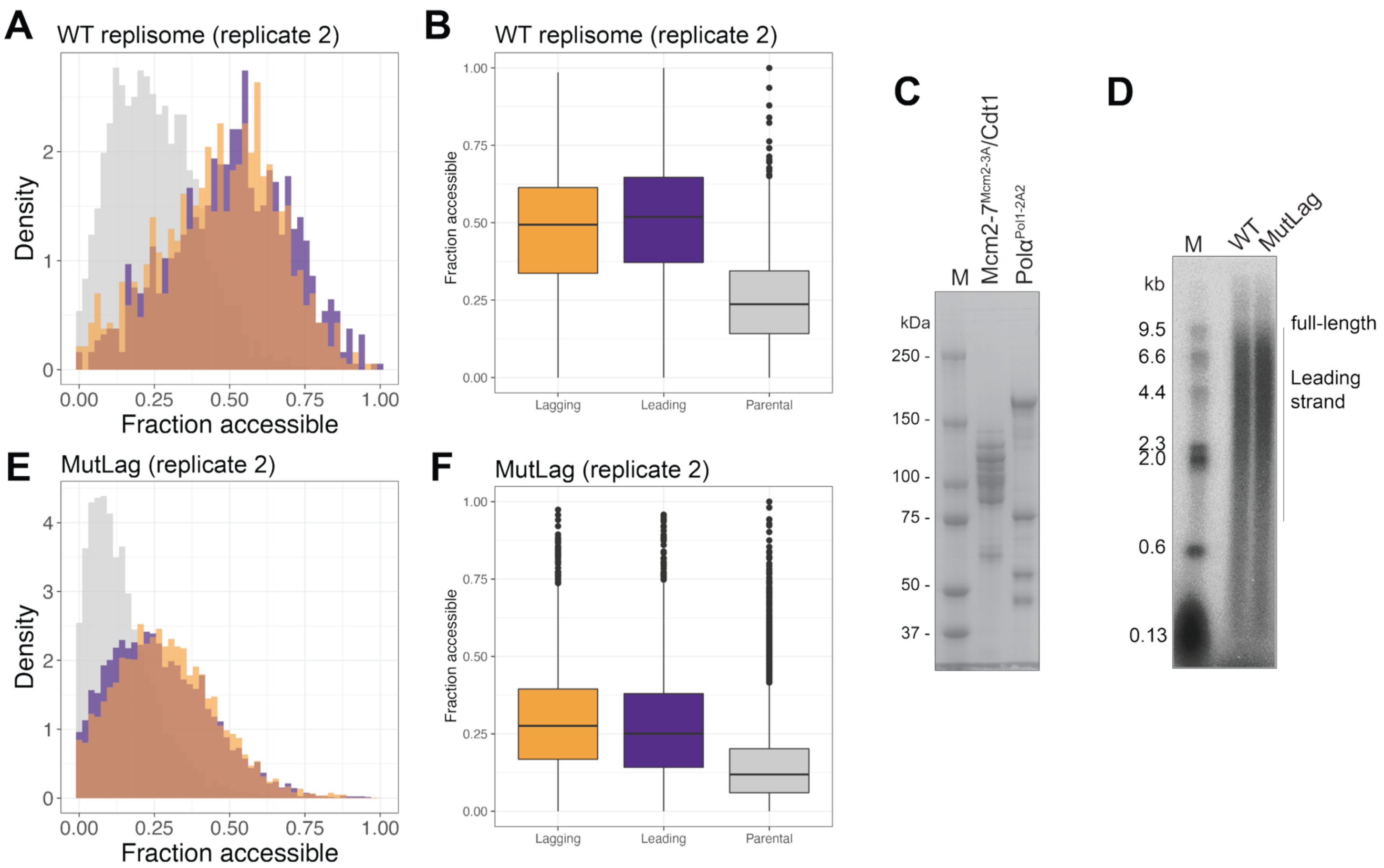


**Supplementary Figure 5: Replicate of mutant lagging-strand replisome confirms increased lagging strand opening for the mutant replisome, while DNA replication remains unaffected. (A)** Accessibility distribution given by the average methylation of single molecules of wildtype replisome replicate 2, separated by non-replicated (grey), lagging strand (orange) and leading strand (purple) classifications. **(B)** Boxplot quantifying the mean accessibility from the distributions as plotted in **(A)** and *p-value* determined by Wilcoxon Rank-Sum test significance level. Wildtype replisome replicate 2 was performed with 20nM of RFC and PCNA. **(C)** SDS PAGE analysis of purified histone chaperone mutant complexes: Mcm2-7/Cdt1 carrying Mcm2-3A (Y79A Y82A Y91A) mutations, and PolA carrying Pol1-2A2 (F58A D62A) mutations. **(D)** Autoradiography of denaturing agarose gel of replication reactions carried out with wildtype or mutant complexes shown in (**C**). No differences in replication efficiency are found when using these mutants. **(E)** Accessibility distribution given by the average methylation of single molecules of mutant lagging-strand replisome replicate 2, separated by non-replicated (grey), lagging strand (orange) and leading strand (purple) classifications. **(F)** Boxplot quantifying the mean accessibility from the distributions as plotted in **(E)** and *p-value* determined by Wilcoxon Rank-Sum test significance level.

Supplementary Figure 6

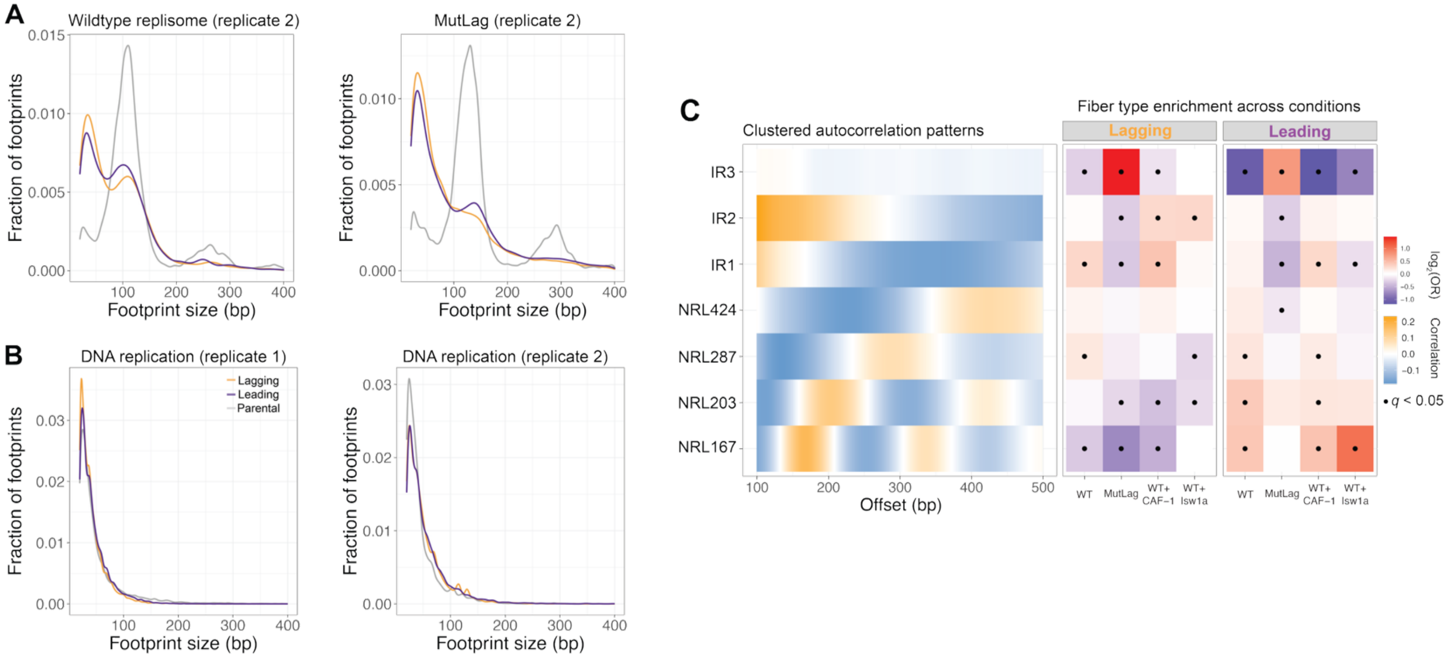


**Supplementary Figure 6: (A) Replicated chromatin is organized differently on the leading- and lagging-strands.** Footprint size distributions plotted as kernel density estimates for wildtype replisome replicate 2 and mutant lagging-strand replisome replicate 2. **(B)** Footprint size distributions plotted as kernel density estimates for naked DNA replication replicate 1 (20 nM RFC and PCNA) and naked DNA replication replicate 2 (50 nM RFC and 200 nM PCNA). **(C) Left:** Single-molecule autocorrelogram using Leiden clustering of four reconstitutions: wildtype replisome, mutant lagging-strand replisome, wildtype replisome + CAF-1 alone and wildtype replisome + ISW1a alone, displaying 7 fiber types characteristic of these reconstitutions. The fiber types are manually annotated as regular, with specific nucleosome repeat lengths (NRLs), or irregular (IR), where either a weak / no autocorrelogram peak was observed. **Right:** Fiber type enrichment determined by Fisher's exact test, to characterize fiber type representation across those different reconstitutions for leading- and lagging-strands.

Supplementary Figure 7

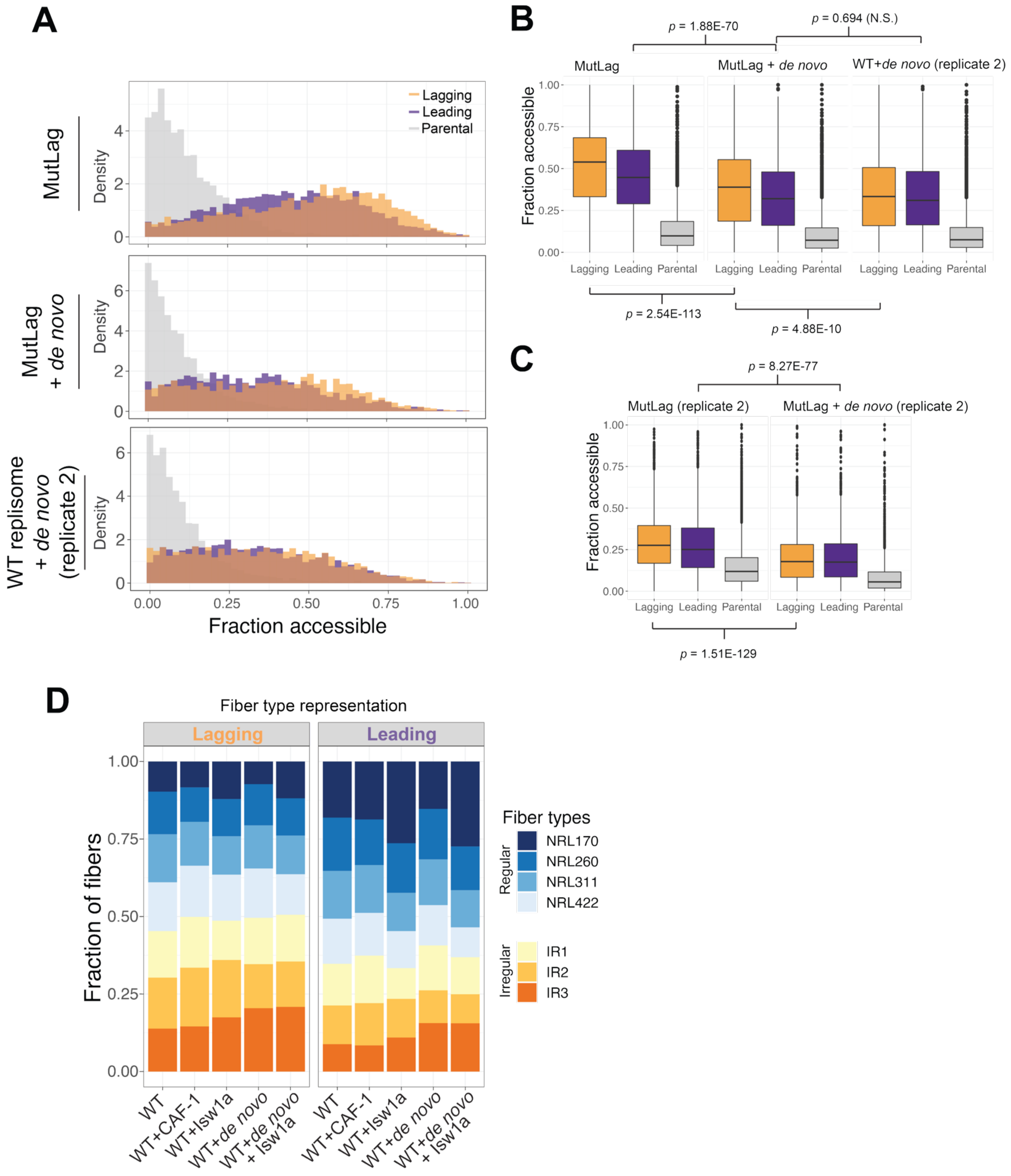


**Supplementary Figure 7: The *de novo* chromatin assembly- and chromatin remodeling- pathways are not sufficient to overcome differences in chromatin organization between both chromatids(A)** Accessibility distribution given by the average methylation of single molecules of replicate 2 of mutant lagging-strand (Mcm2–

3A and Pol1-2A2) replisome reconstitution with or without *de novo* histone deposition, and replicate 2 of WT replisome+*de novo* histone deposition separated by non-replicated (grey), lagging strand (orange) and leading strand (purple) classifications. **(B)** Boxplot quantifying the mean accessibility from accessibility distributions and *p-values* as determined by Wilcoxon Rank-Sum test significance level for mutant lagging-strand replisome, mutant lagging-strand replisome + *de novo* and wildtype replisome + *de novo*. **(C)** Boxplot quantifying the mean accessibility from accessibility distributions and *p-values* as determined by Wilcoxon Rank-Sum test significance level for replicates 2 of mutant lagging-strand replisome and mutant lagging-strand replisome + *de novo*. **(D)** Relative quantification of fiber type representation per reconstitution for leading and lagging strands. Fiber types were obtained from single-molecule autocorrelogram using Leiden clustering of five reconstitutions: wildtype replisome, wildtype replisome + CAF-1 alone, wildtype replisome + ISW1a alone, wildtype replisome + *de novo*, wildtype replisome + *de novo* + ISW1a, where all *de novo* conditions include soluble histones.

### Materials and Methods

#### Protein expression and purification

ORC, cdc6, Mcm2/7-cdt1, S-CDK, DDK, Dpb11, GINS, cdc45, Mcm10, RPA, Polα-prim, Polε, Ctf4, Sld3/7, Sld2 and TopoII were expressed and purified as in (*21*). Polδ, Csm3/Tof1, Topo I, RFC and PCNA were purified as described in (*22*). Mrc1 was expressed and purified following the procedure described in (*48*). Nhp6 and ISW1A were expressed and purified following (*24*). Nap1 was also expressed and purified following (*24*), with a slight change of buffers where KXPO4 pH 7.6 was replaced by 20 mM Tris-HCl pH 7.2 in all buffers. Asf1 was also expressed and purified as described in (*24*), with the dialysis step replaced by a Superdex S200 Increase column, and the protein was eluted in buffer containing 30 mM HEPES pH 7.6, 100 mM KCl, 10% glycerol, 5 mM DTT and 0.1% IGEPAL CA-630. CAF-1 was expressed and purified as in (*49*). Mutants Mcm2-7$^{Mcm2\text{-}2A}$/Cdt1 (containing Mcm2-2A: Y79A Y82A Y91A) and Polα$^{Pol1\text{-}2A2}$-prim (containing Pol1-2A2 F58A D62A) were cloned using PCR mutagenesis and were purified following the same protocols as the wild-type proteins. Lyophilized individual *S. cerevisiae* histones were purchased from the Histone Source at CSU, Fort Collins, CO, USA. Histones H3, H4, H2A and H2B were refolded as octamers and stored in 2M NaCl containing buffer as described in (*50*). Additional protein purification protocols are:

*FACT*: Spt16 and Pob3 subunits were co-overexpressed under the bidirectional *GAL1-10* promoter in a yJF1 background. The transformed cells were obtained from Dr. John Diffley (yAE89, unpublished). Cells were grown overnight at 30°C in 2% raffinose until cell density reached 2-5x$10^7$ cells/mL. Protein expression was induced by addition of 2% galactose, and continued for 3 hours. The cells were frozen drop-wise in liquid nitrogen and crushed in a freezer mill. Cell powder was washed in lysis buffer (20 mM Tris-HCl pH 8.0, 500 mM NaCl + 0.3 mM PMSF and protease inhibitor tablet, cOmplete, Roche), followed by centrifugation at 32,000 rpm, 1hour at 4°C. The cleared supernatant was loaded to 1.5 mL washed anti-Flag M2 beads and incubated for 1 hour at 4°C. On a disposable 20-mL column (BioRad Econo-Pac), the protein-bound beads were washed in 100 CV wash buffer (20 mM Tris-HCl pH 8.0, 200 mM NaCl, 10% glycerol, 1 mM DTT). The protein was eluted in 1 CV wash buffer + 0.5 mg/mL 3x FLAG peptides, followed by 2 CV wash buffer + 0.25 mg/mL 3x FLAG peptides and 2 CV wash buffer. The eluate was pooled, concentrated in an Amicon Ultra-4 Centrifugal Filter Unit 30 MWCO and loaded on a Superdex S200 Increase 10/300 GL column equilibrated in GF buffer (20 mM Tris-HCl pH 8.0, 200 mM NaCl, 10% glycerol, 1 mM DTT). The fractions of interest were pooled, concentrated and stored at -80°C.

*Fen1*: pRS303-CBP-TEV-Fen1 plasmid received from Dr. John Diffley was overexpressed under the *GAL1-10* promoter in yJF1 cells. Cells were grown overnight at 30°C in 2% raffinose. When cells reached 2-5x$10^7$ cells/mL, 2% galactose was added to induce protein expression for 3 hours. Cells were lysed in a freezer mill. Cell lysate powder was resuspended in lysis buffer (25 mM Tris-HCl pH 7.5, 150 mM NaCl, 10% glycerol, 0.01% IGEPAL CA-630, 1 mM EDTA, 0.3 mM PMSF and protease inhibitor tablet, cOmplete, Roche) and cell debris were removed by centrifugation at 32,000 rpm for 1 hour at 4°C. The supernatant was loaded on Sepharose 4B calmodulin beads (GE Healthcare) pre-washed in binding buffer (25 mM Tris-HCl pH 7.5, 150 mM NaCl, 10% glycerol, 0.01% IGEPAL CA-630, 2 mM $CaCl_2$, 1 mM DTT). Protein was bound to the beads for 1.5 hours at 4°C, and subsequently loaded to a disposable 20 mL column (BioRad Econo-Pac). Beads were washed with 100 CV binding buffer, and protein was eluted in 1 mL fractions elution buffer (25 mM Tris-HCl pH 7.5, 150 mM NaCl, 10% glycerol, 0.01% IGEPAL CA-630, 2 mM EDTA, 2 mM EGTA, 1 mM DTT). Protein containing fractions were pooled and loaded on a 1-mL HiTrap Heraparin HP column (Cytiva). Protein was eluted over a gradient of Heparin buffer (25 mM Tris-HCl pH 7.5, 10% glycerol, 0.01% IGEPAL CA-630, 1 mM DTT) A (150 mM NaCl) to B (1M NaCl). Fractions containing Fen1 were pooled and dialyzed for 2 hours against dialysis buffer (25 mM Tris-HCl pH 7.5, 100 mM NaCl, 10% glycerol, 0.01% IGEPAL CA-630, 1 mM DTT). The protein was concentrated and stored at -80°C.

*Cdc9*: yJY33 cells (unpublished) received from Dr. John Diffley (Gal-cdc9-2xFlag) were grown overnight in 2% raffinose at 30°C. When cells reached cell density between 2-5x$10^7$ cells/mL, protein expression was induced with 2% galactose for 2 hours. Cell lysate derived from freezer mill was resuspended in cdc9 lysis buffer: 25 mM Tris-

HCl pH 7.5, 150 mM NaCl, 10% glycerol, 0.01% IGEPAL CA-630, 1 mM DTT, 0.3 mM PMSF and protease inhibitor tablet, cOmplete, Roche. Insoluble material was cleared via centrifugation at 32,000 rpm for 1 hour at 4°C. The supernatant was incubated with 1.5 mL anti-Flag M2 affinity resin (Sigma) for 2 hours at 4°C. The resin was loaded on a disposable 20 mL column (BioRad Econo-Pac) and subsequently washed with 100 CV cdc9 lysis buffer. Cdc9 was eluted in 1 CV lysis buffer + 0.5 mg/mL 3x FLAG peptides, followed by 2 CV lysis buffer + 0.25 mg/mL 3x FLAG peptides and 2 CV lysis buffer. The eluate was pooled, and applied to a 5 mL HiTrap Q HP (Cytiva). Protein elution was done over a gradient from in buffer (25 mM Tris-HCl pH 7.5, 10% glycerol, 0.01% IGEPAL CA-630, 1 mM DTT) from 150 mM NaCl to 800 mM NaCl. The protein was concentrated in an Amicon Ultra-4 Centrifugal Filter Unit 30 MWCO and loaded on a Superdex S200 Increase 10/300 GL (Cytiva) column equilibrated in GF buffer (25 mM Tris-HCl pH 7.5, 150 mM NaCl, 10% glycerol, 0.01% IGEPAL CA-630, 0.01% IGEPAL CA-630, 1 mM DTT). The fractions containing the clean protein were concentrated and stored at -80°C.

*Pif1*: pJHH16 (6XHis tagged Pif1 cloned into pET28a) plasmid was received from Dr. John Diffley. The plasmid was transformed into BL21 Rosetta cells (homemade competent). Cells were grown in LB under selection with 100 μg/mL ampicillin and 34 μg/mL chloramphenicol at 37°C to an $OD_{600}$ of 0.5. Protein expression was induced with 0.1 mM IPTG for 20 hours at 16°C. Cell pellet was washed in 40 mM KXPO4 pH 7.0, 100 mM NaCl, 8 mM MgCl2, 0.01% IGEPAL CA-630, 1 mM DTT, 10% glycerol and protease inhibitor tablet, cOmplete, Roche. Cell lysis was done via sonication, followed by supernatant clearing via centrifugation at 35000 xg for 30 min at 4°C. The supernatant was incubated with Ni+ sepharose beads (prepared from Chelating Sepharose Fast Flow resin, Cytiva) for 2 hours at 4°C. The resin was collected in a 20 mL disposable column (BioRad Econo-Pac) and washed extensively with 40 mM KXPO4 pH 7.0, 300 mM NaCl, 8 mM MgCl2, 0.01% IGEPAL CA-630, 1 mM DTT, 10% glycerol + 2 mM ATP + 20 mM imidazole. Elution of Pif1 was done in 10x 1 mL fractions in 40 mM KXPO4 pH 7.0, 150 mM NaCl, 8 mM MgCl2, 0.01% IGEPAL CA-630, 1 mM DTT, 10% glycerol + 200 mM imidazole. Pooled elutions were diluted 2-fold in 40 mM KXPO4 pH 7.0, 150 mM NaCl, 8 mM MgCl2, 0.01% IGEPAL CA-630, 1 mM DTT, 10% glycerol and loaded on a HiTrap SP HP (Cytiva). Pif1 was eluted in 40 mM KXPO4 pH 7.0, 8 mM MgCl2, 0.01% IGEPAL CA-630, 1 mM DTT, 10% glycerol in a gradient from 150 mM NaCl to 1 M NaCl. Peak clean fractions were pooled, concentrated in an Amicon Ultra-4 Centrifugal Filter Unit 30 MWCO and applied to a Superdex 200 Increase 10/300 GL column (Cytiva), and eluted in 40 mM KXPO4 pH 7.0, 150 mM NaCl, 8 mM MgCl2, 0.01% IGEPAL CA-630, 1 mM DTT, 10% glycerol + 0.5 mM EDTA. Peak fractions were pooled, concentrated, aliquoted and stored at -80°C.

**Replication templates**

Circular plasmid vJY22 (*22*), a gift from Dr. John Diffley, is an ARS1-containing 10.6 Kb plasmid and was used as a template for our replication reactions. The plasmid was used as naked or chromatinized templates. Biochemical assays were performed using circular plasmid, as well as MscI-digested linear DNA. All of the reactions intended for sequencing were performed using MscI-linearized DNA, in its naked or chromatinized version.

Plasmid digestion was done using 10 μg DNA and 7.5 μL MscI in 1X CutSmart Buffer (New England Biolabs) and a final reaction volume of 120 μL for 1.5 h at 37°C. If full digestion was not confirmed on agarose gel, the DNA was subjected to another round of digestion after enzyme heat inactivation at 80°C for 20 min, and gradient cool down to 37°C. In this second round of digestion, the reaction was supplemented with 7.5 μL MscI enzyme, and the reaction was conducted for 1.5 h at 37 °C. In the event of a second round of digestion, heat inactivation of MscI was only performed in between the two steps of digestion. Subsequently, the linear DNA was purified using phenol-chloroform extraction, and the purified DNA pellet resuspended in TE buffer.

**Chromatin assembly template preparations**

Chromatin arrays were produced both on circular and linear templates, following a similar protocol as originally developed in (*51*) and adapted in (*22*, *23*). Briefly, Nap1 (3 μM), ISW1a (30 nM) and yeast histone octamer (370 nM) were incubated on ice for 10 min, in chromatin assembly buffer (25 mM Hepes-KOH pH 7.6, 100 mM KOAc, 10 mM $Mg(OAc)_2$, 0.01% IGEPAL CA-630, 5% glycerol, 0.1 mg/mL BSA, 3 mM ATP, 40 mM creatine phosphate and 0.28 mg/mL creatine phosphate kinase). DNA template (3nM) was then added to the protein mix, and chromatin assembly started for 10 min at 30ºC. After 10 min, ORC was added (20 nM) and chromatin assembly continued for 50 min. Chromatin templates were then purified by applying 50 μL reaction to Microspin S-400 HR columns (Cytiva) equilibrated in 3x 400 μL 25 mM Hepes-KOH pH 7.6, 100 mM KOAc, 0.02% IGEPAL CA-630, 10 mM $Mg(OAc)_2$, 1 mM DTT.

***In vitro* DNA and chromatin replication reactions**

Reaction volumes for the biochemical reconstitutions ranged between 30 – 80 μL. Reactions used for SAMOSA footprinting and PacBio library preparation had a final volume of 270 μL. Independent on the final volume, all reactions were run as below.

Mcm2-7 helicase loading step was done in one third of the final volume in 25 mM Hepes-KOH pH 7.6, 100 mM K-glutamate (naked DNA) or 100 mM KOAc (chromatin), 10 mM $Mg(OAc)_2$, 0.01% IGEPAL CA-630, 1 mM DTT, 5 mM ATP, 0.1 mg/mL BSA and 20 – 22.5 nM ORC, 45 nM cdc6, 40 – 100 nM Mcm2-7/Cdt1 and 5 nM (naked) or ~ 3 nM (purified chromatin) template. The reaction was conducted for 30 min at 30ºC. Phosphorylation reaction was then initiated by addition of 50 nM DDK at 30ºC for another 30 min. Finally, replication started by adding first FF500 buffer (final concentrations: 17 mM Hepes-KOH pH 7.6, 167 mM, 7 mM $Mg(OAc)_2$, 0.007% IGEPAL CA-630, 0.7 mM DTT, 2 mM ATP, 0.07 mg/mL BSA, 50 μM dATP-dCTP-dGTP-dTTP (dTTP 100% replaced by BrdUTP in sequencing reactions), 22 nM α-[32P]-dCTP (biochemical experiments only) and 130 μM CTP-GTP-UTP). Then, replication protein mix were was added at final concentrations as follows: 40 nM cdc45, 30 nM Dpb11, 20 nM S-CDK, 5 nM Mcm10, 20 nM TopoII, 10 nM TopoI, 20 nM Polε, 20 nM Polα, 20 nM Ctf4, 25 nM Sld3/7, 50 nM Sld2, 210 nM GINS, 100 nM RPA, 20 nM Csm3/Tof1, 20 nM Mrc1, 20-200 nM PCNA, 20-50 nM RFC, 10 nM Polδ, 40 nM Fen1, 40 nM cdc9, 5 nM Pif1. Chromatin replication reactions included 80 nM FACT and 400 nM Nhp6 added in the replication proteins mix. Reactions containing the *de novo* pathway included 40 nM CAF-1, 70 nM Asf1 and 30 nM yeast histone octamer, added from a separate mix immediately after replication reaction initiation by addition of the replication proteins mix. Replication reactions were conducted for 40 min at 30ºC. Replication was stopped by adding a final concentration of 25 mM EDTA. MNase assays for a biochemical chromatin readout were done on 30-40 μL reaction volume, not quenched by EDTA. The samples were diluted 1.7x to reduce salt concentration, followed by addition of 5 mM $CaCl_2$ and 370 U MNase (NEB) in 30 uL reaction. MNase digestion at 37ºC was conducted for 5 min and quenched by addition of 50 mM EDTA.

**Biochemical experiments gel readout**

To readout DNA replication, samples were cleaned-up for unincorporated nucleotides using MicroSpin G-50 columns (Cytiva). Samples were then denatured in 100 mM NaOH, 2% sucrose, bromocresol green as loading dye and 15-25 μL samples were run on 0.7% alkaline agarose gels for 18 h at 45 V. DNA was precipitated on gel following 2 cycles of 15 min in ice cold 5% TCA with TCA refreshment. The gel was dried in 2× chromatography Whatman paper in a gel dryer for 2 h at 60°C. Gel was exposed to a phosphor screen for 1-4 days and imaged using Amersham Typhoon Biomolecular Image.

To readout Okazaki fragment maturation and replication termination, 50 μL replication samples quenched by EDTA were first treated with 0.1% SDS and 3.5 μL proteinase K at 42ºC for 30 minutes. DNA was cleaned up following phenol-chloroform extraction and resuspended in 30 μL TE. The DNA was then digested with 1 μL NheI enzyme in 50 μL reaction containing 1x CutSmart buffer at 37ºC for 1 hour. 25 μL digested DNA was loaded on a 0.8% agarose gel in 1x TAE at 20 V overnight. The gel was dried in between 2× chromatography Whatman paper in a gel dryer for 2 h at 60°C. Gel was exposed to a phosphor screen for 1-4 days and imaged using Amersham Typhoon Biomolecular Image.

Biochemical chromatin samples obtained from MNase reactions were treated with 10 μL Proteinase K at 42ºC for at least 30 minutes, up to an hour. Samples were then cleaned up using MinElute PCR purification kit (Qiagen) and eluted in 10 μL TE. Samples were loaded on a native 1.3% agarose gel in 1x TAE and gel ran for 3 hours at 180 V (30 cm gels). Gel was stained in SYBr Gold for 1 hour to show chromatin assembly on total DNA, followed by treatment using 2 cycles of 15 min ice cold 5% TCA. The gel was then dried in between 2× chromatography Whatman paper in a gel dryer for 2 h at 60°C. Gel was exposed to a phosphor screen for 1-4 days and imaged using Amersham Typhoon Biomolecular Image.

**Calculation of gel-based efficiencies of replication**

To calculate gel-based efficiencies of replication, we used an approach similar to (*32*). Briefly, a 1:120 dilution sample was taken from the original reaction and spots of 0.5 µL were made in triplicate on filter paper beside the gel. An autoradiography scan of the gel and spots was taken and ImageQuant TL software was used to assess the counts in the spots and the replicated samples. The dCTP concentration in the reaction is 53 µM. The maximum amount of DNA that can theoretically be synthesized per microliter of reaction is 53 µM × 1 µL × 330 g/mol × 4 nucleotides, or about 70 ng. If this much DNA is synthesized, all the radioactive dCTP will be incorporated in the DNA. To calculate the actual amount of DNA synthesized, the number of counts in 0.5 µL of replicated sample is divided by the total counts in the 0.5 µL of the spot. Then this value is multiplied by 70 ng, to obtain the actual amount of dCTP incorporated in the newly replicated DNA. Finally, the percentage of replicated DNA, or the efficiency of replication, is obtained by dividing the amount of incorporated dCTP calculated in the previous step, divided by the amount of input DNA.

**SAMOSA on *in vitro* (chromatin) templates and replicated reactions**

SAMOSA methylatransferase footprinting reactions were performed on templates: naked DNA and chromatin arrays (inputs) and replicated reactions on naked DNA and chromatin. The nonspecific adenine EcoGII methyltransferase (New England Biolabs, high concentration stock 2.5 × 104 U/mL) was used as described in (*31*). 1350 ng inputs (naked orchromatin) were methylated using 1 µL EcoGII in reactions at a final volume of 200 µL and containing 1× CutSmart Buffer and 1 mM S-adenosyl-methionine (SAM, New England Biolabs) and incubated at 37 °C for 30 min. 1 µL SAM was spiked in after 15 min. The entire volume of EDTA-quenched replication reactions (308.6 µL – 1750 ng for naked DNA and estimated ~ 500 ng for chromatin after Microspin S-400 HR column) was methylated with 2-3.5 µL EcoGII in 500 µL containing 1× CutSmart Buffer and 1 mM S-adenosyl-methionine (SAM, New England Biolabs) and incubated at 37 °C for 30 min. 2 µL SAM was spiked in after 15 min.

To purify inputs treated with EcoGII, 0.45% SDS and 2.5 µL proteinase K (NEB, 800 U/mL) were added and incubated at 65°C for 2 hours. To purify replicated samples treated with EcoGII, 0.45% SDS and 6.25 µL proteinase K (NEB, 800 U/mL) were added and incubated at 65°C for 2 hours. Next, samples were cleaned up using 2x SPRI beads volume. DNA binding to the beads was conducted at 37°C for 30 minutes with interval mixing. Beads were placed on a magnetic rack and 2x washed with 80% ethanol, allowing 1 minute to remove salt content. Spun beads briefly to collect excess ethanol. Allowed quick drying before resuspending beads in 25 µL elution buffer (10 mM Tris, pH 8.5, 0.1 mM EDTA) and incubating at 37°C for 30 minutes with interval mixing to elute DNA. DNA concentration was measured using Qubit High Sensitivity DNA assay (Invitrogen).

**Pacbio long-read sequencing**

Purified footprinted DNA from *in vitro* replication reactions, between 200 – 500 ng per reaction, was used as input for half-reactions of the Pacbio SMRTbell prep kit 3.0. Briefly the kit steps are: end repair and DNA repair, SMRTbell circular adapter ligation with unique barcodes, nuclease to remove unligated DNA, and lastly, clean-up using 1X SMRTbell beads. Final library concentration was measured using Qubit High Sensitivity DNA Assay. Libraries were then pooled and sequenced on PacBio Sequel II 8M SMRTcells at 120 – 150 pM loading concentration per cell using the Sequel II Binding Kit 3.2 and 30 hour movie time following 2 hour pre-extension time.

**Data processing and analysis**

All custom scripts for the steps detailed below are available at https://github.com/RamaniLab/repli-SAMOSA-ChAAT.

**Long-read sequencing preprocessing**

Preprocessing steps were performed using a custom shell script with Pacbio packages. Briefly, the steps are: *ccs* (v6.99.99) processes subreads into circular consensus reads, *lima* (v2.13.0) demultiplexes barcoded samples, and *pbmm2* (v1.17.0) aligns the CCS (circular consensus sequencing) reads to the reference fasta file. This produced aligned CCS BAM files which were used for all subsequent processing.

**Single-molecule chromatin accessibility**

Adenine methylation by EcoGII was detected using the IPD (interpulse duration) from Pacbio polymerase kinetics as previously described (*31*, *32*). Briefly, unmethylated and fully methylated template DNA was sequenced and used to train a truncated singular value decomposition model (SVD) with 40 components to reduce representation. Next a neural network (NN) model was trained to predict the mean log IPD at each potentially methylated base in the unmethylated control, which allowed us to subtract that prediction from the measured value to get residuals. A two-component simple mixture model was then fit to the residual values for the unmethylated and methylated templates to determine a cutoff for calling a base a methylated, and filters were applied to remove bases without well-defined cutoffs or at GATC contexts. Next, a Viterbi HMM (hidden Markov model) was applied to predicted methylation calls to construct a most likely state path for each read, which is stored in a pickled dictionary with ZMW (zero-mode waveguide) number as the key.

**Strand-specific BrdU detection at the origin of replication**

As above, a NN model was trained per base, in both forward and reverse directions, to get residuals at each position. Reads for clustering were filtered to only those that fully cover the region around the origin (+/- 1000 bp) based on the BAM alignment. The residuals for T bases, where BrdUTP would be incorporated during replication, on the Watson (reverse T) and Crick (forward A) strands, were concatenated and NaN values filled with zeros, with a maximum of 10 NaNs tolerated. The concatenated residuals per ZMW were then clustered with the Leiden algorithm at a resolution of 0.2 with graph structure defined using the metric of correlation with ten nearest neighbors, clusters were filtered to those that represent a minimum of 0.5% of the total clustered data to remove a long tail of extremely small clusters. The mean residual values were plotted and observed to find clusters of asymmetric residual values, where either Watson or Crick values had low variability and the other strand had high variability, frequently reaching a mean residual of over 0.5. The strand with high variability in residuals, meaning the polymerase frequently stalled or sped up, indicates that strand has BrdUTP incorporation. Using these clustering results, each clustered ZMW per sample was assigned Watson, Crick, or No BrdU status in a metadata file used for downstream analyses.

**Downstream data analyses**

*Plotting residuals per cluster*

For all clusters from residual clustering above, residuals were pulled based on ZMW and stored in a matrix where each column represents a base and then means per base were calculated, using *numpy.nanmean*. The resulting data were plotted in R for the two most abundant non-replicated clusters and the Watson and Crick replicated clusters, looking at T positions within 1 kb of the origin.

*Calculating footprint sizes and positions*

Inaccessible stretches from the HMM output were called as footprints with length and midpoint position per ZMW as previously described in (*31*).

*Tabulating footprint frequency by size*

Footprints calculated above were stratified by size (short < 80 bp; nucleosomal 80 – 160 bp; oligonucleosomal 160 + bp). Nucleosomal footprints per ZMW was used to determine the number of nucleosomes per molecule. Footprint counts per stratification per midpoint position within +/- 1kb of the origin were tabulated and plotted.

*Footprint position reproducibility*

Short footprints between 20 and 80 bp within 1 kb of the origin on BrdUTP$^{(+)}$ molecules were selected from two independent replicates of replicated DNA. Footprint midpoints were binned (bins = 300) and plotted, then

Spearman and Pearson correlation statistics for frequency of footprints per bin between the replicates was calculated using *scipy.stats*.

*Defining leading and lagging strands*

For all replicated molecules, 2000 bp were selected, centered on the ORC binding site beginning at 8410 bp. Data were then split at this center; for the right-hand side, 8410 – 9409 bp, Watson-replicated strands were defined as leading and Crick-replicated strands as lagging. For the left-hand side, 7410 – 8409 bp, Watson-replicated strands were defined as lagging and Crick-replicated strands as leading. For subsequent analyses using leading and lagging terminology, the right and left-hand data were combined. Parental strands are the same window for BrdUTP$^{(-)}$ molecules.

*Calculating fraction accessible*

Fraction accessible was calculated for the entire molecule or for the leading and lagging strand windows depending on the figure. In all cases, the mean of the HMM values per molecule for the given window was taken using *numpy.nanmean*.

*Autocorrelation spacing analysis*

Autocorrelation was calculated in Python using *ccf* (Cross-Correlation Function), as previously described in *(31, 32, 38, 52)*, for the first 1000 bp for rightward and leftward moving forks. Autocorrelation results for all chromatin samples were then clustered with the Leiden algorithm (*37*) at a resolution of 0.7 with graph structure defined using the metric of correlation with 15 nearest neighbors. Clusters representing 90% of the total data were retained. Peaks within the autocorrelation were annotated using *scipy.find_peaks* to define NRL (nucleosome repeat length) and clusters without clear peaks were defined as IR (irregular).

*Leading and lagging strand fiber type bias*

Using the cluster annotations (fiber types) from the autocorrelation analysis above, we used Fisher's Exact Tests (*scipy.stats.fishers_exact*) to compute enrichment of each cluster per sample and per strand. Using the $\log_2$(OR) we then computed relative enrichment of fiber types across all reconstitutions and secondly computed strand bias of fiber types per reconstitution. A Fisher's $q$ value of $< 0.1$ or $<0.05$ was noted as significant on enrichment plots, as described in each figure legend.
